# Leveraging peptide substrate libraries to design inhibitors of bacterial Lon protease

**DOI:** 10.1101/689877

**Authors:** Brett M. Babin, Paulina Kasperkiewicz, Tomasz Janiszewski, Euna Yoo, Marcin Drąg, Matthew Bogyo

## Abstract

Lon is a widely-conserved housekeeping protease found in all domains of life. Bacterial Lon is involved in the recovery from various types of stress, including tolerance to fluoroquinolone antibiotics, and is linked to pathogenesis in a number of organisms. However, detailed functional studies of Lon have been limited by the lack of selective, cell-permeable inhibitors. Here we describe the use of positional scanning libraries of hybrid peptide substrates to profile the primary sequence specificity of bacterial Lon. In addition to identifying optimal natural amino acid binding preferences, we identified several non-natural residues that were leveraged to develop optimal peptide substrates as well as a potent peptidic boronic acid inhibitor of Lon. Treatment of *E. coli* with this inhibitor promotes UV-induced filamentation and reduces tolerance to ciprofloxacin, phenocopying established *lon*-deletion phenotypes. It is also non-toxic to mammalian cells due to its increased selectivity for Lon over the proteasome. Our results provide new insight into the primary substrate specificity of Lon and identify substrates and an inhibitor that will serve as useful tools for dissecting the diverse cellular functions of Lon.

## Introduction

Lon is a widely-conserved housekeeping protease, found in bacteria, archaea, and eukaryotic mitochondria and chloroplasts^1^. All Lon orthologs feature a AAA+ ATPase domain that unfolds protein substrates and a proteolytic domain that catalyzes the hydrolysis of those substrates^2^. The importance of bacterial Lon has been determined mostly through studies using *Escherichia coli lon* mutants and via biochemical analyses of recombinant enzyme. Lon has myriad regulatory functions related to stress-response^3,4^, including roles in the SOS response to DNA damage^5^, defense against reactive oxygen species^6^, heat shock^7^, amino acid starvation^8^, and phage integration^9^. Phenotypic consequences of *lon* deletion include the inability to recover normally from UV-induced DNA damage and the reduced persistence of *lon* mutants following fluoroquinolone treatment^10,11^. In the context of pathogenesis, *lon* mutants of many bacteria are defective for infection. These include *Pseudomonas aeruginosa* in lung infection models of mice and rats^12^, *Salmonella enterica* in macrophages and systemic infection of mice^13^, and *Brucella abortis* in macrophages and spleen infections of mice^14^.

Due to its roles in stress-response, Lon is an interesting target for small-molecule inhibition. A selective inhibitor enables dynamic studies of Lon proteolysis in a variety of physiological contexts and, based on the links between Lon and pathogenesis, has the potential to be useful as a therapeutic agent^15^. Furthermore, specific inhibition of Lon protease activity would allow separation of its proteolytic functions from those involving chaperone activity or binding of DNA and polyphosphate. For example, controlled inhibition of Lon would be useful for clarifying the role the protease plays in persistence. While a defect in fluoroquinolone tolerance in *lon* mutants has been established for many years^16^, there has been substantial debate about the mechanism by which Lon contributes to this phenomenon^17,18^. A recently-proposed model for the role of Lon in persistence which involves the degradation of toxin-antitoxin modules has since been disproven^19–21^. The current model involves regulation through degradation of the cell-division inhibitor SulA, the same mechanism by which Lon directs recovery from other sources of DNA damage. According to this model, Lon proteolytic activity would be important primarily when SulA is overexpressed as part of the SOS response. The ability to precisely control Lon inactivation (i.e., by addition of a small-molecule inhibitor) would be critical to test this hypothesis.

While a number of small-molecule Lon inhibitors have been identified, to our knowledge, none have been used to test the consequences of Lon inhibition within live bacterial cells. Lon features a serine-lysine dyad in its active site, notably different from the canonical serine-histidine-aspartic acid found in many serine proteases^22,23^. A likely consequence of its noncanonical active site is that many broad-spectrum serine protease inhibitors have poor activity against the enzyme. Early studies noted that *E. coli* Lon could be inhibited by the serine protease inhibitors diisopropyl fluorophosphate^24^ and dansyl fluoride,^25^ but only at mM concentrations. Inhibitors with slightly greater potency include 3,4-dichloroisocoumarin, other coumarin derivatives^26^, and oleanane triterpenoids^27^. Another class of Lon inhibitors comprise peptidic compounds that couple an amino acid recognition moiety with an electrophilic ‘warhead’ that covalently reacts with the active-site serine to inactivate the enzyme. Examples of inhibitors with activity for Lon include Z-Gly-Leu-Phe-chloromethylketone^28^, as well as the human proteasome inhibitors MG132^29, 30^ (featuring an aldehyde warhead), and MG262 and bortezomib (**BZ**)^31, 32^ (featuring boronic acid warheads). In addition, a larger, hexapeptide boronic acid inhibitor of Lon was generated from the amino acid sequence of the natural λN Lon substrate^33^. These peptidic inhibitors take advantage of amino acid sequences that are tolerated by Lon, but none have been optimized for the enzyme, nor counter-screened for potential off-target binding or inhibition.

One of the most significant issues for current Lon inhibitors is their high level of cross reactivity with the proteasome. This leads to significant toxicity, making them ineffective as tools to study Lon function in cells. It is striking that many proteasome inhibitors have cross-reactivity with Lon, considering the differences in the active-sites: hydrolysis by the proteasome is catalyzed by an N-terminal threonine. Crystal structures of *Meiothermus taiwanensis* Lon revealed that the boronic acid warheads of MG262 and **BZ** bind covalently to the active-site serine, like their covalent modification of the threonine hydroxyl in the proteasome^32^. This strong covalent reactivity of boronates towards active site hydroxyls explains the dual potency for Lon and the proteasome.

We set out to develop selective inhibitors that could be used to specifically block Lon protease activity in cells. We hypothesized that the identification of highly selective peptide substrates could be leveraged to generate an optimized inhibitor using established electrophilic warheads. This strategy builds on a body of work from our groups and others describing the conversion of peptide substrates to inhibitors and activity-based probes for diverse protease targets^34^, including caspases^35^, cathepsins^36^, human neutrophil elastase^37^, human neutrophil serine protease 4^38^, both the human^39^ and *Plasmodium*^40, 41^ proteasome, and proteases important in *Mycobacterium tuberculosis* pathogensis^42^ and Zika virus infection^43^. By screening a large combinatorial library of peptide substrates, we identified a sequence of amino acids optimized for Lon. Based on this screening data, we designed a peptidic boronic acid inhibitor with potent activity for Lon and reduced potency for the human 20S proteasome. This compound was non-toxic to mammalian macrophages and is able to phenocopy classic *lon* deletion phenotypes in *E. coli*. We expect this compound to serve as a tool for studying the role of Lon-mediated proteolysis during stress response and pathogenesis.

## Results and Discussion

The primary sequence specificity of Lon has been examined using individual fluorogenic peptide substrates^28^ as well as by identifying the cleavage sites for a number of endogenous Lon substrates^44–46^. However, there has not yet been a comprehensive and unbiased profiling of its amino acid preferences. We therefore performed a screen for fluorogenic peptide substrates using a hybrid combinatorial substrate library (HyCoSuL) that has been successfully applied to other protease targets^37,39,47^. Lon was purified after recombinant expression in *E. coli* (Figure S1). We chose to use libraries of tetrapeptides in which the P1 residue directly adjacent to the site of hydrolysis was fixed as a phenylalanine in order to ensure recognition by Lon. These libraries are made up of a set of sub-libraries in which 121 natural and non-natural amino acids are scanned through each of the P2 and P3 positions on the substrate (Figure 1a). Cleavage by Lon of each sub-library containing a fixed P2 or P3 residue is used to determine the overall specificity patterns at those positions. Initial analysis of the natural amino acid libraries provides some insight into the potential cleavage sites of native protein substrates. We found that at both the P2 and P3 positions, multiple residues are accepted, suggesting an overall broad specificity of the protease (Figure 1b). At each position, bulky or hydrophobic residues yielded the best substrates. This result is consistent with previous reports of favored peptide substrates that feature Ala, Leu, and Phe residues, and various analyses of the cleavage sites within protein substrates. In addition, these results support the model in which Lon is involved in degrading unfolded hydrophobic domains of endogenous substrates. It should be noted that, while information about peptide substrate preferences may be useful for determining preferred cleavage sites and cleavage rates of natural substrates, data from our peptide library screens cannot be used to determine preferences for protein substrates that are dictated by interactions with other domains of the enzyme (e.g., the substrate recognition domain).

**Figure 1.**
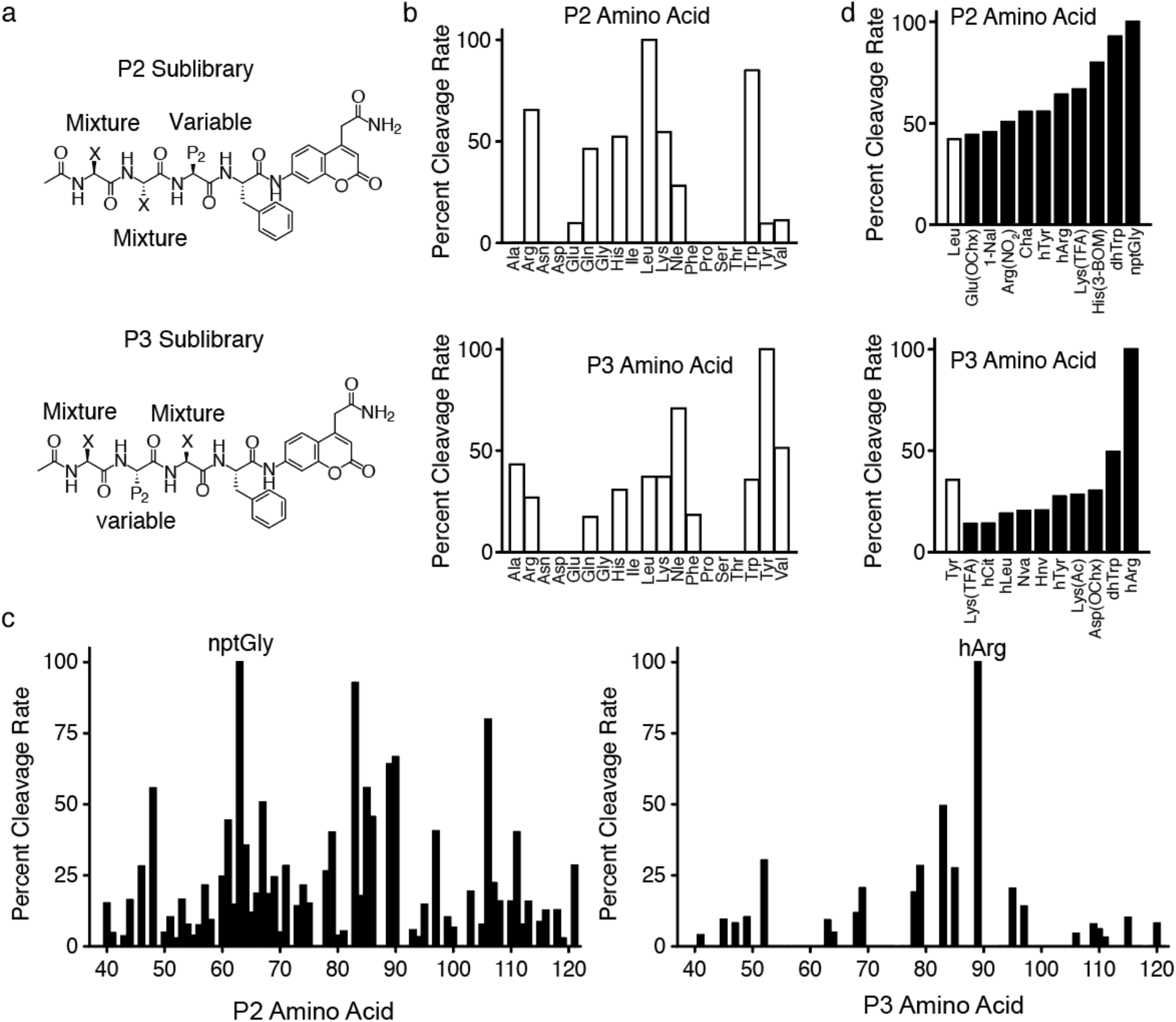
HyCoSuL screening of Lon substrates. Structures of (a) P2 and P3 fluorogenic HyCoSuL libraies. Positions comprising equimolar mixtures of 18 natural amino acids and norleucine (Nle) as a substitute for Met and Cys (not included in the library) are designated by an X. The remaining variable position is held constant as the indicated natural or non-natural amino acid for each sub-library. Plot of the relative cleavage rates for (b) natural amino acids in the P2 (top) or P3 (bottom) positions and (c) non-natural amino acids in the P2 (left) or P3 (right) positions. (d) Plots of cleavage rates of the best non-natural amino acids for the P2 (top) or P3 (bottom) positions. The best natural amino acid (white bar) at each position is included for comparison. Results for each sub-library were normalized to the amino acid with the fastest cleavage rate, indicated by an arrow, and data represent the mean of two independent screens. Amino acid type is indicated by white (natural) or black (non-natural) bars. D-amino acids (Nos. 20-36) exhibited no cleavage and were excluded from the plot in (c). Amino acid abbreviations and structures are as reported in Kasperkiewicz, et al^37^. The list of amino acids and normalized cleavage rates can be found in Table S1.

We next performed substrate cleavage analysis using the libraries containing non-natural amino acids to get a broader perspective on the substrate specificity of Lon (Figure 1c-d; Table S1). Interestingly, Lon accepted a diverse array of non-natural amino acids at the P2 position, with more than 50% of the library exhibiting measurable cleavage. In contrast, it was more stringent at the P3 position and showed a strong preference for a single non-natural amino acid, L-homoarginine (hArg). For both positions, the most-preferred amino acids contained bulky side chains. To verify the results of the combinatorial library screening, we generated a set of fluorogenic tri-peptide substrates that contained the newly identified P3 hArg as well as the fixed P1 Phe and a morpholine acetate N-terminal cap. We then varied the P2 position using amino acids selected from the best substrates identified in the substrate screen (Figure 2a). For comparison, we used Mo-Leu-Leu-Phe-ACC (**1**), a peptide substrate containing only natural amino acids. This substrate yielded kinetic parameters nearly identical to those previously reported for the Lon substrate, Glt-Ala-Ala-Phe-MNA^28,48^. In contrast, all of the substrates containing the P3 hArg greatly outperformed **1**, with specificity constants (k_cat_/K_M_) as much as 17-fold higher for the best substrate, **5**, which contains a neopentylglycine (nptGly) at the P2 position (Figure 2b, Figure S2a).

**Figure 2.**
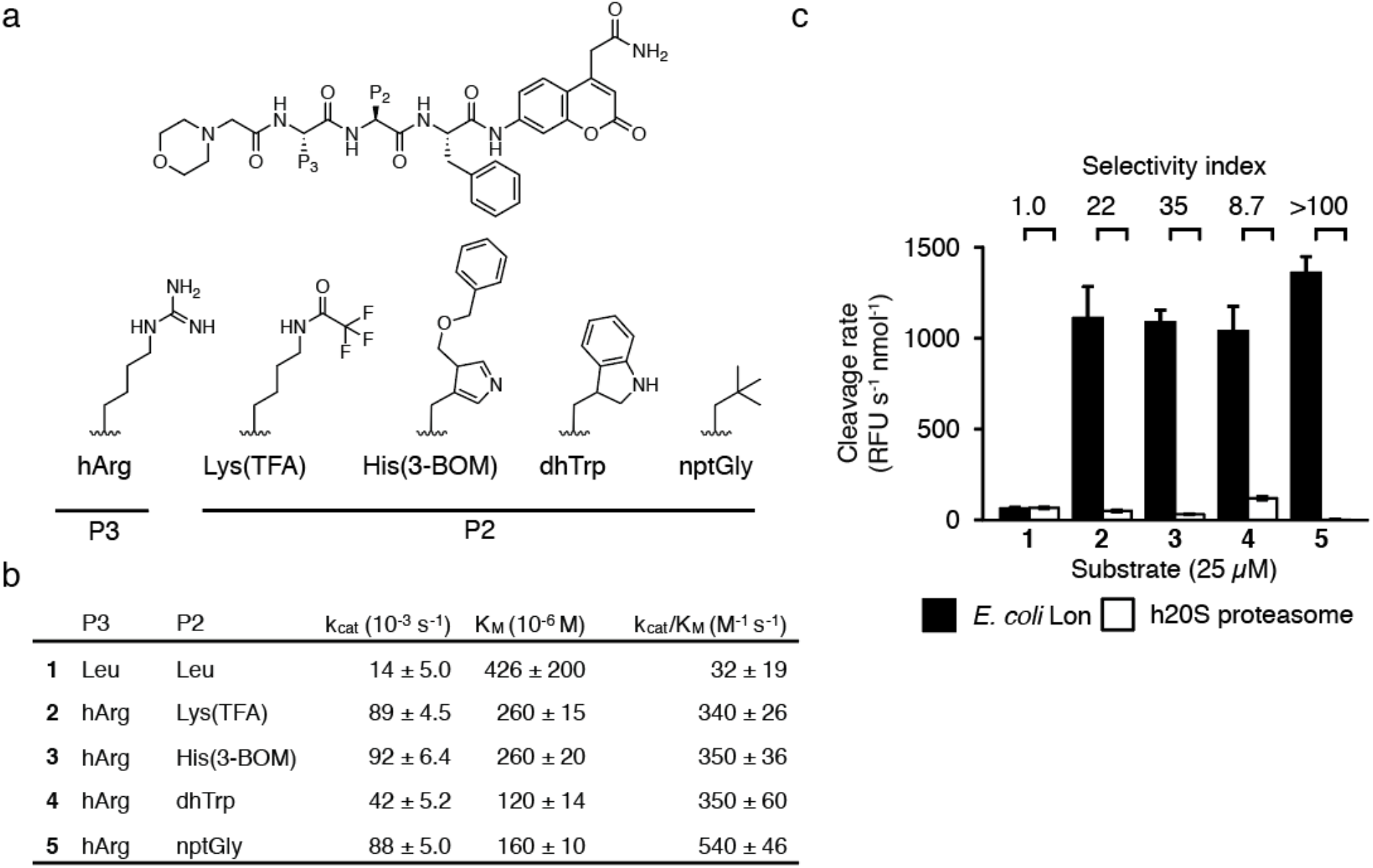
Design of selective Lon substrates. (a) General structure of fluorogenic Lon substrates. Amino acids used in the P3 and P2 positions are shown. (b) Kinetic parameters for cleavage by Lon for each substrate (mean ± standard deviation, n=3). (c) Cleavage rates for each substrate by Lon (black) and h20S (white) (mean ± standard deviation, n=3). Rates were normalized to total enzyme amount. Selectivity values were calculated as the ratio of cleavage rates. Cleavage rate of 25 μM **5** by h20S was negligible. Raw kinetic data are shown in Figure S2.

Having determined optimal substrates for Lon, we set out to use these scaffolds to build a potent, covalent inhibitor of Lon. The fact that several classes of covalent inhibitors have been reported suggests that the choice of electrophile is important for the optimal inhibitor design. We therefore screened our existing focused library of electrophilic protease inhibitors^49^ to identify an appropriate electrophile. This set of compounds includes diverse, reactive moieties that form permanent covalent bonds with active-site serine, threonine, or cysteine residues, including diphenyl phosphonates, vinyl sulfones, epoxy ketones, chloroisocoumarins, vinyl ketones, and triazole ureas. To screen this set of ~1,200 compounds we established an *in vitro* enzyme assay using our optimized fluorogenic peptide substrate **5**. Our initial screen at a high concentration (10 μM) of the compounds identified a small number of hits that abolished Lon activity (Figure S3a-b). While we identified hits within all warhead classes, even the most potent compounds from the screen had IC_50_ values well above that of the human proteasome inhibitor **BZ**, which has previously been reported as an inhibitor of Lon (Figure S3c). We therefore decided to focus on using the reversible covalent boronic acid electrophile in **BZ** to make an optimized Lon inhibitor.

We suspected that converting any one of the Lon substrates (**2**-**5**) to a boronic acid would yield a potent inhibitor. Because peptide boronic acids have been shown to be highly effective inhibitors of the human proteasome, counter screening for proteasome inhibition is essential to avoid high toxicity due to this cross-reactivity. To identify the peptide scaffold that would likely yield the most selective Lon inhibitor, we evaluated cleavage of the substrates by both Lon and the human 20S proteasome (h20S). These results showed that the non-optimized substrate **1** containing the Leu-Leu-Phe sequence was cleaved equally effectively by both Lon and the proteasome while substrates containing the optimized P3 hArg (**2-5**) were primarily cleaved by Lon and not the human proteasome. In fact, cleavage of **2-5** by the proteasome was so weak that it did not saturate and as a result, we were unable to determine kinetic parameters for those substrates (Figure S2b-c). In lieu of kinetic constants, we compared normalized cleavage rates for a fixed substrate concentration (Figure 2c). These results confirmed that substrates **2**-**5** were selective for Lon, with **5** being the most selective. This substrate showed essentially no detectable cleavage by the h20S. This result is consistent with a HyCoSuL screen of h20S that showed that peptides featuring hArg in the P3 position are poor substrates for the β1 and β5 subunits^39^.

To leverage the identified substrate specificity of Lon into the design of a selective inhibitor, we synthesized a hybrid compound containing the P2 and P3 residues of substrate **5** combined with the P1 Leu and N-terminal pyrazinamide cap of **BZ** to generate Pyz-hArg-nptGly-Leu-B(OH)_2_ (**6**, Figure 3a). Both **6** and **BZ** exhibited potent, time-dependent inhibition of recombinant Lon (Figure S4a-b), with IC_50_ values after 60 min of inhibitor pre-incubation approaching the active-site concentration used in the assay (Figure S4c), suggesting covalent inhibition. Kinetic analyses (Figure 3b-c, Figure S4d-e) showed **6** to be a more potent Lon inhibitor than **BZ** with a two-fold higher k_inact_/K_I_ driven primarily by improved potency (i.e., a lower K_I_ value) (Figure 3d). To test for activity toward the human proteasome, we pre-treated purified h20S with each compound and then labeled subunit active sites with the fluorescent, activity-based probe MV151 (Figure 3e)^50^. Competition for active-site labeling of β1 and β5 subunits of h20S required a 10-fold higher concentration of **6** than **BZ** (50 vs 5 μM). Inhibition assays using fluorogenic peptides specific for each subunit similarly showed an increase in IC_50_ values for **6** compared to **BZ** for the β1 and β5 subunits (Figure S4f-g). Surprisingly, we also saw some inhibition of the β2 subunit by **6**, despite the strong preference of this “trypsin-like” subunit for Arg at the P1 position. Together these results confirm that the increase in Lon potency of **6** compared to **BZ** was accompanied by a reduction in binding to the β1 and β5 subunits of the proteasome. More importantly, the Lon inhibitor **6** was not cytotoxic to murine macrophages at doses as high as 10 μM. This is in stark contrast to **BZ** which kills the same cells with an EC_50_ of 160 nM (Figure 3f). Thus, the drop in potency of **6** towards the proteasome is sufficient to eliminate the toxicity in mammalian cells and suggests that it should be a valuable new compound for use in cell biological studies of Lon function.

**Figure 3.**
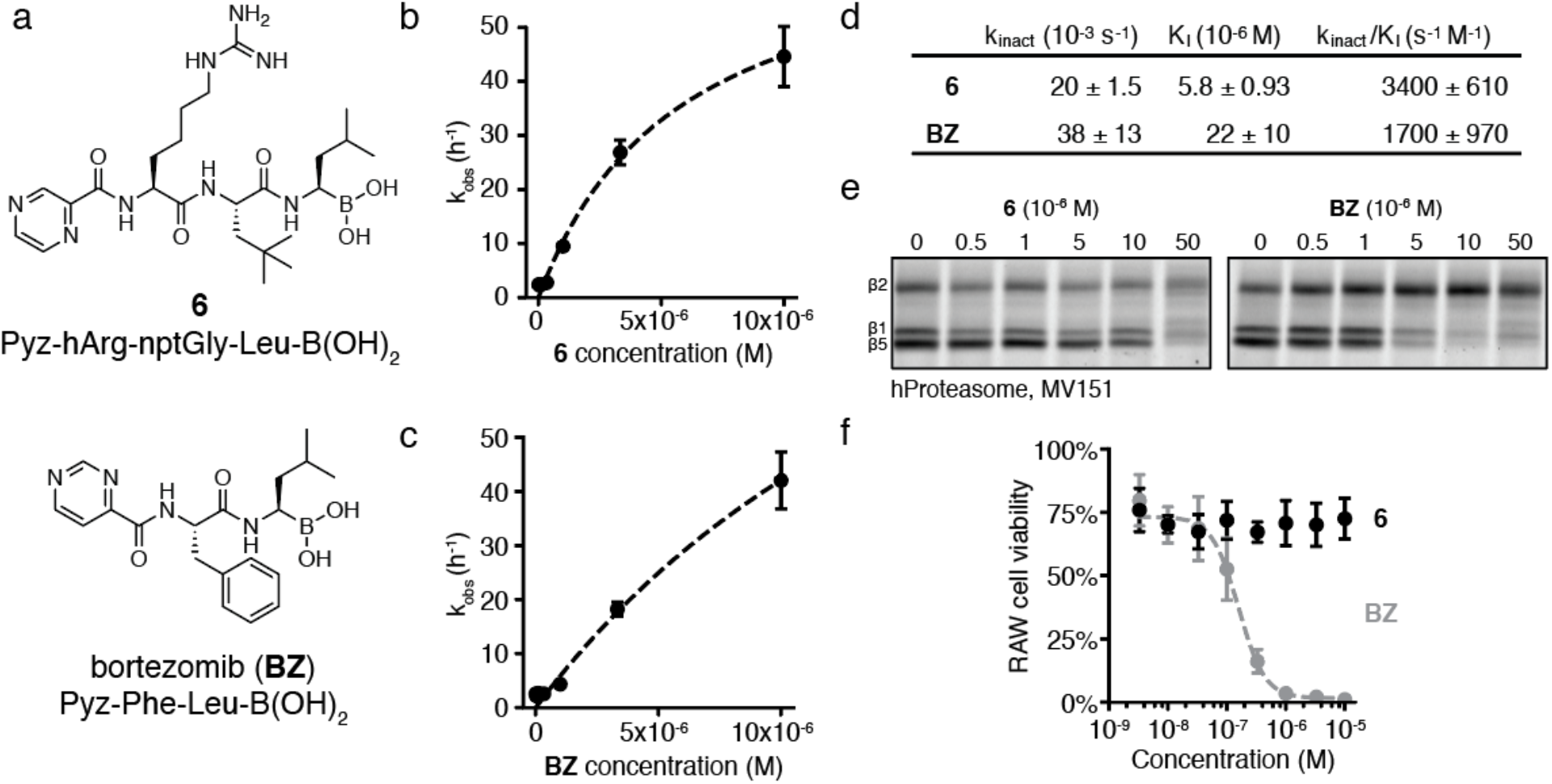
Design of a selective Lon inhibitor. (a) Structures of the designed Lon inhibitor **6** and **BZ**. Kinetic analysis of time-dependent inhibition of Lon *in vitro* by (b) **6** and (c) **BZ** (mean ± standard deviation, n=3). Dashed line indicates the fit used to calculate kinetic parameters. Raw kinetic data are shown in Figure S4. (d) Kinetic parameters of time-dependent inhibition of Lon (mean ± standard deviation, n=3). (e) h20S treated with **6** or **BZ**, then labeled with MV151. Proteins were separated by SDS-PAGE and scanned for MV151 fluorescence. Images are representative of two independent experiments. (f) Viability of murine RAW macrophages following 24 h treatment with **6** (black) or **BZ** (gray). Dashed line indicates the fit used to obtain the IC_50_ for **BZ**. Viability was quantified by normalizing CellTiter-Blue fluorescence to that of untreated cells (mean ± standard deviation, n=3).

Although Lon plays important roles in stress response and pathogenesis, *lon* is a nonessential gene and deletion mutants grow normally in the absence of exogenous stress. We generated a clean-deletion of *lon* (Figure S5a-b) and found that neither genetic disruption of *lon* nor treatment with 100 μM **6** or **BZ** had an effect on exponential growth rates (Figure S5c). One of the first observed consequences of *lon* mutation in *E. coli* was the filamentation of cells after UV-induced DNA damage^51^. DNA damage causes upregulation of the cell-division inhibitor SulA as part of the SOS response. Lon-mediated degradation of SulA allows cells to resume division after recovery from stress. In the absence of Lon, SulA concentrations remain high and cells grow but cannot divide, resulting in extended filaments. We hypothesized that if **6** was a selective inhibitor of Lon then treatment of *E. coli* should phenocopy the eponymous “long” filamentation phenotype found in *lon* cells. As expected, outgrowth following UV-stress resulted in long filaments in the *lon* deletion strain but not in wild-type or *sulA* mutant cells (Figure 4a). Treatment with **6** during outgrowth following UV-stress led to a dose-dependent increase in filamentation (Figure 4b, Figure S6a). In a *sulA* mutant strain, **6** had no effect on filamentation, similar to observations of *lon sulA* double mutants^52^. Quantification of cell area for more than 500 cells per condition showed increases in the maximum cell area and in the percent of cells that were filamented (i.e., with area greater than 4 μm^2^, Figure 4c).

**Figure 4.**
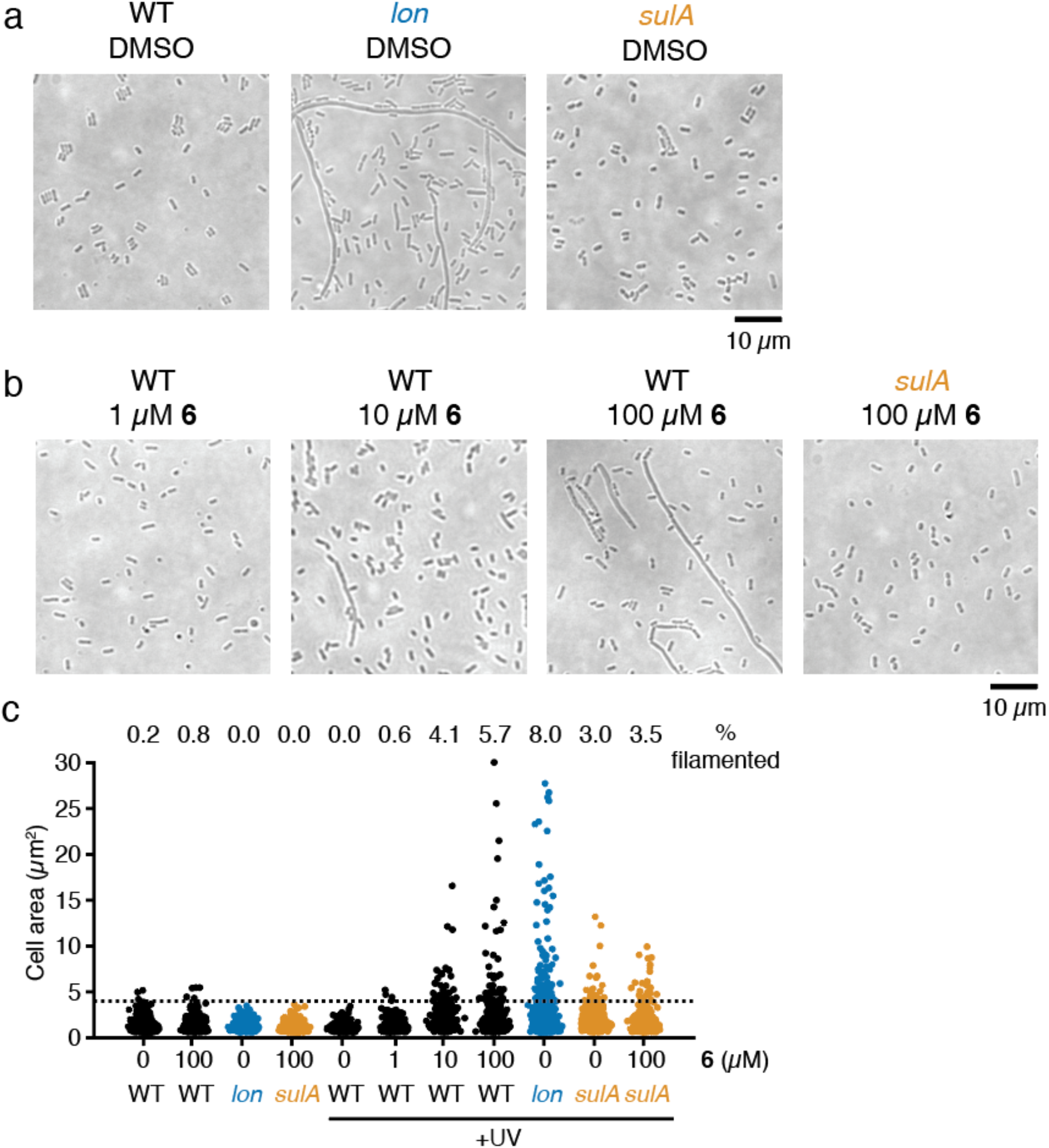
Lon inhibition induces *E. coli* filamentation following UV stress. (a-b) Representative phase contrast images of *E. coli* strains exposed to UV light and then diluted into LB containing (a) DMSO or (b) various concentrations of **6**. Cultures were grown for 6 h after UV-exposure and then imaged. Images are representative of two independent experiments. (c) Quantification of cell area for cells without UV-treatment or cells treated as in (a-b). The percent of cells with area greater than 4 μm^2^ (dashed line) is indicated (n>500 for each condition). Additional images are presented in Figure S6a.

Lon is also implicated in recovery from DNA damage caused by fluoroquinolone antibiotics^16,53^. Most cells treated with such antibiotics die, but a small subpopulation (typically 0.01% of the initial population) tolerate antibiotic exposure and can replicate after removal of the antibiotic. So-called persister cells^54^ are reduced in a *lon* knockout strain. Like UV-induced filamentation, Lon’s role in persistence depends on the presence of SulA, with *lon sulA* double mutant strains producing a similar number of persisters as wild-type cells^10,17,55^. We therefore predicted that co-treatment of cells with **6** and ciprofloxacin would reduce persister cell formation. Neither *lon* deletion nor treatment of wild-type cells with 100 μm **6** altered the overall MIC of ciprofloxacin (0.0125 μg/ml). However, compared to wild type, we consistently observed a statistically-significant reduction in the fraction of cells that tolerated ciprofloxacin for both the *lon* mutant strain and wild-type cells treated with **6** in both rich (LB, Figure 5a) and minimal media (M9, Figure S6b). Importantly, this effect was abrogated in the *sulA* mutant strain, suggesting it results from inhibition of Lon. Furthermore, the effect was time-dependent, with both *lon* and **6**-treated cells exhibiting faster death than wild type (Figure 5b). The effect was also concentration-dependent, with the extent of effect from **6** treatment matching that of the *lon* knockout strain at high concentrations (Figure 5c).

**Figure 5.**
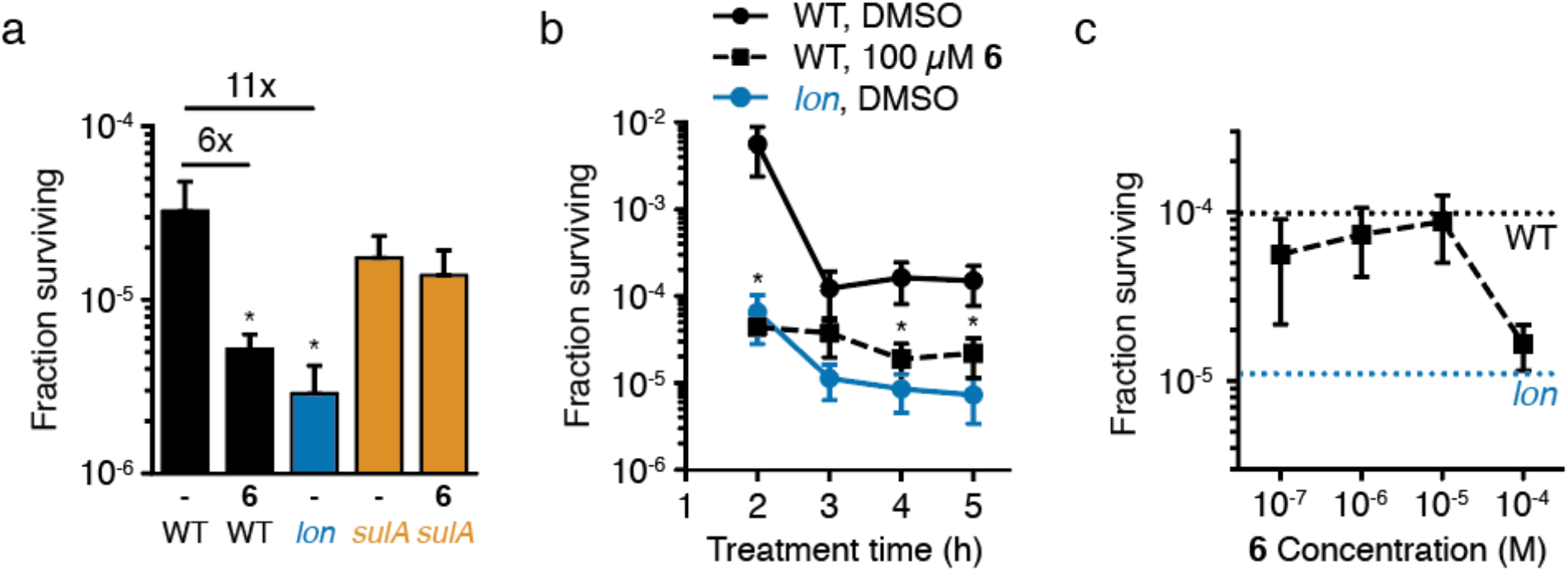
Lon inhibition decreases ciprofloxacin persister cells. All panels show the fraction of *E. coli* cells surviving treatment with 10 μg/mL ciprofloxacin in LB. (a) Cells were co-treated with ciprofloxacin and either DMSO (-) or 100 μM **6** for 4 h (mean ± standard deviation, n=3; unpaired t-tests comparing each sample to wild-type cells treated with DMSO: *, p<0.05). Fold reduction compared to wild-type treated with DMSO is indicated above. (b) Time course of killing for cells co-treated with ciprofloxacin and DMSO or 100 μM **6** (mean ± standard deviation, n=3; unpaired t-tests comparing each sample to wild-type cells treated with DMSO at the same time point: *, p<0.05; *lon* was significantly different at all time points). (c) Dose-dependent effect of **6** in reducing cell survival. Cells were co-treated with ciprofloxacin and the indicated dose of **6** for 4 h (mean ± standard deviation, n=3; unpaired t-tests comparing each sample to wild-type cells treated with DMSO: p=0.054 for 100 μM **6**). Dashed lines indicate the fraction surviving of wild-type or *lon* cells treated with DMSO. For all experiments, cultures were centrifuged, washed, and cell numbers were determined by spot dilution plating on LB agar. Panel (a) is representative of four independent experiments.

The SulA-dependent effects of **6** on UV-induced filamentation and ciprofloxacin persister formation strongly suggest that the compound inhibits Lon in cells. Incomplete phenocopying and the requirement for a high dose (e.g., 100 μM for cellular effects) are likely due to some combination of active efflux of the compound and permeability barriers. Both of these issues are common challenges for treating gram-negative bacteria with small molecules^56^. Encouragingly, there is evidence that other boronic acid inhibitors can enter *E. coli* cells^57, 58^, so we expect that modifications to increase the permeability of **6** will lead to further improved potency against live cells.

We leveraged amino acid preferences of Lon to develop both an improved fluorogenic substrate and a boronic acid inhibitor of Lon with increased selectivity over the proteasome. Our substrate screening results build on previous observations that Lon prefers to cleave peptides with bulky, hydrophobic residues, consistent with its role in degrading denatured proteins during stress responses. In our initial screen for inhibitors, Lon was poorly inhibited by electrophiles such as diphenyl phosphonates and chloroisocoumarins, which are potent inhibitors of many proteases with canonical serine-histidine-aspartic acid catalytic triads. This observation, along with the potency of proteasome inhibitors toward Lon, highlight the unusual nature of the serine-lysine dyad in its active site. Structural analyses of Lon inhibition by **6** would confirm the hypothesized covalent interaction with the active-site serine and would help to explain the structural basis for Lon’s preferences for bulky amino acids and the role that hArg plays in enhancing substrate and inhibitor binding. In the future, novel Lon inhibitors may be identified by exploring alternative warheads such as β-lactams^59^, or nitriles^60^ which have activity toward serine-lysine dyads in signal peptidases and the UmuD family of proteases.

We expect **6** to be a useful compound for studying the roles that Lon plays in stress-response and pathogenesis. The use of a small molecule inhibitor rather than genetic disruptions (e.g., *lon* deletion or active-site mutants) introduces a level of dynamic flexibility to studies of Lon. Additionally, it provides a means to disentangle Lon’s proteolytic activity from other functions of the multidomain complex, such as ATPase activity and its ability to bind and respond to DNA. In our cellular experiments, we observed **6**-mediated effects on cellular physiology both when the compound was added during recovery from (outgrowth after UV exposure) or concurrent with (co-treatment with ciprofloxacin) stress. These observations suggest that Lon inhibition during or after stress has similar effects, at least for the SulA-mediated models of stress response tested here. Our data also show that **6** is not toxic to macrophages, meaning it can be used to test inhibition of bacterial Lon in cell culture models of infection and pathogenesis. Finally, because it is a covalent inhibitor, it can be converted to a fluorescent or otherwise affinity-labeled probe in order to visualize Lon activity within living cells. This compound should therefore greatly expand the scope of future studies of Lon function.

## Methods

Methods for Lon purification, inhibitor screens, determination of substrate and inhibitor parameters, generation of the *lon* knockout strain, and chemical syntheses are described in the Supporting Information.

### HyCoSuL screens

HyCoSuL screens were performed in Corning opaque 96-well plates. Each well contained 99 μl of Lon in assay buffer (250 mM Tris, pH 8.0, 1 M KCl, 100 mM MgCl_2_, and 1 mM ATP). Lon was added to a final hexamer concentration of 190 nM (P2 library) or 570 nM (P3 library). HyCoSuL substrates were added to a final concentration of 100 μM and kinetic fluorescence measurements (ex. 355 nm, em. 460 nm) were taken at 37 °C for at least 30 min starting immediately after substrate addition (Spectramax Gemini XPS, Molecular Devices). The substrate hydrolysis rate (RFU/s) was calculated from the linear portion of each progress curve. The amino acid with the highest cleavage rate was set to 100%, and remaining amino acids were adjusted accordingly. Each library was screened twice and results are presented as mean values.

### Kinetic analysis of substrates

Lon and h20S substrate cleavage assays were performed in black 96- or 384-well plates. For Lon experiments, each well contained 25 μl 2X Lon assay buffer, 0.5 μL of 100 mM ATP (1 mM final concentration), and 40 nM final concentration of Lon hexamer. For ATP regeneration, 0.75 μl of 5 mg/ml creatine kinase (75 μg/ml final concentration) and 4 μl of 50 mM creatine phosphate (4 mM final concentration) were included. Water was added to a final volume of 40 μl. For h20S experiments, each well contained 25 μl 2X h20S buffer (100 mM Tris, pH 7.5, 200 mM NaCl), 1 mM DTT, 2 nM final concentration of h20S (BostonBiochem), 24 nM final concentration of PA28 (BostonBiochem), and water to a final volume of 40 μl. To begin the reaction, 10 μl of each substrate was added from a 5X stock, and fluorescence (ex. 360 nm, em. 460 nm) was measured every minute for 1 h at 37 °C in a microplate reader (BioTek Cytation 3).

### Kinetic analysis of inhibitors

Inhibition assays were performed under the same conditions as for substrate kinetics. Compounds were added from a 100X stock in DMSO (0.5 μl). For pre-incubation, compounds were added to the enzyme mixture in each well and plates were incubated at 37 °C for the indicated time. For experiments without pre-incubation, compounds were added to the working stock of substrate. Substrates (10 μl of 250 μM working stock) were added to the enzyme mixture and fluorescence (ex. 360 nm, em. 460 nm) was measured every minute for 1 h at 37 °C in a microplate reader (BioTek Cytation 3). For Lon, **5** was the substrate. For h20S, Z-LLE-AMC, Boc-LRR-AMC, and Suc-LLVY-AMC were substrates specific for the β1, β2, and β5 subunits, respectively. Proteasome substrates were purchased from BostonBiochem.

### Proteasome labeling

For each compound, 1 μl of 20X stock in DMSO was added to a sample of h20S (10 nM) in 19 μl labeling buffer (50 mM Tris, pH 7.5, 5 mM MgCl_2_, 1 mM DTT) and incubated for 1 h at 37 °C. To label proteasome subunits, 0.5 μl of 80 μM MV151 (final concentration 2 μM) was added and incubated for an additional 2 h at 37 °C. Labeling was quenched by addition of 4X Laemmli sample buffer, samples were incubated for 5 min at 95 °C, and samples were separated by SDS-PAGE. MV151 fluorescence was imaged using a Typhoon 9410 Imager on the Cy3 channel (Amersham Biosciences).

### RAW cell viability

RAW 264.7 murine macrophages were cultured in DMEM with 4.5 g/l glucose, 4 mM L-glutamine, and 10% FBS (Invitrogen) at 37 °C with 5% CO_2_. Cells were split and seeded into a 96 well-plate to 5×10^3^ cells per well with 50 μl of medium. To each well was added 49 μl of medium with 1 μl of 100X compound in DMSO (1% final DMSO concentration). Cells were incubated with compound for 24 h then treated with 20 μl CellTiter-Blue (Promega) for 4 h. Cell viability was quantified by measuring fluorescence in a microplate reader (BioTek Cytation 3). Fluorescence values were normalized to untreated cells. Incubation with 1% DMSO reduced cell viability compared to untreated cells, but the effect was independent of compound or dose.

### Bacterial strains and growth conditions

Bacteria were cultured with shaking in LB (Fisher), 2xYT (Teknova), or M9 at 37°C, unless otherwise indicated. M9 contained 6 g/l Na_2_HPO_4_, 3 g/l KH_2_PO_4_, 1 g/l NH_4_Cl, 0.5 g/l NaCl, 0.5% (w/v) glucose, 1 mM MgSO_4_, 0.1 mM CaCl_2_, and 0.34 mg/l thiamine HCl. The *lon* mutant strain was generated by clean deletion of the coding region of *lon* using homologous recombination with CRISPR-Cas9 selection^61^. The *sulA* mutant strain (*sulA773(del)::kan*) was obtained from the Keio Collection^62^. Growth rates were determined by measuring OD_600_ of 100 μL cultures grown at 37°C in a 96-well plate in a microplate reader overnight.

### UV treatment and microscopy

Overnight cultures of wild-type, *lon*, or *sulA* strains were grown in LB. Cultures were diluted to OD_600_ 0.1 in LB and grown for 1 h. Cells were pelleted by centrifugation (8,000 g for 5 min), resuspended in 0.1 volume of 10 mM MgSO_4_ and transferred to glass tubes. Cells were irradiated with 900 J/cm^2^ 254 nm light (Stratagene Stratalinker 2400). Control cultures were resuspended in MgSO_4_ as above, but were not irradiated. Cells were diluted 1:25 into LB with compound added from 100X stock in DMSO (1% final DMSO concentration) and grown for 6 h at 37 °C with shaking in the dark. For imaging, 4 μl of each culture was applied to 2% agarose pads^63^. Phase contrast microscopy was performed on a Zeiss LSM700 confocal microscope with a Plan-Apochromat 63x/1.4 objective. Twenty-five images were captured via tile scan for each condition. Quantification of cell area was performed with the MicrobeJ^64^ plugin for ImageJ. Regions of interest containing at least 500 cells were analyzed using default settings for bacterial detection. A minimum cell area of 0.9 μm^2^ was used to exclude non-cellular debris.

### Ciprofloxacin treatment and persister cell quantification

For MIC measurements, overnight cultures of wild-type or *lon* strains were grown in LB, then diluted 1:50 into Mueller Hinton Broth 2 (Sigma). Diluted cultures (50 μl) were aliquoted into wells in a 96-well plate, each containing 50 μl medium and 2X the final concentration of ciprofloxacin. For persister experiments, overnight cultures of wild-type, *lon*, or *sulA* strains were grown in LB or M9. Cultures were diluted to OD_600_ 0.01 in the same medium and incubated for 2 h. Cultures were treated with 10 μg/ml ciprofloxacin (Sigma) from a 100X stock in water and compound from a 100X stock in DMSO (1% final DMSO concentration). Aliquots (100 μl) of each culture were removed at the indicated time, pelleted by centrifugation (8,000 g for 5 min), washed once with PBS, and resuspended in 100 μl PBS. Cells were serially diluted in PBS, 10 μL spots were spotted onto LB agar plates, and plates were incubated for 16-24 h at 37 °C. Colonies were counted to determine CFU.

### Software

Statistical analysis, fitting, and plotting were performed with Python v. 3.6.0, Scipy v. 1.1.0, Numpy v. 1.13.3, Matplotlib v. 3.0.3, and Seaborn v. 0.9.0. Microscopy data were analyzed in ImageJ. DNA sequence analysis was performed in SnapGene 4.3.10. Figures were assembled in Adobe Illustrator CS6.

## Acknowledgments

The authors appreciate critical feedback of the manuscript from Markus Lakemeyer.

## Funding sources

B.M.B. was supported by the Stanford Dean’s Fellowship and the A. P. Giannini Foundation. The work was funded in part by grants from the National Institutes of Health (R01EB026332, R01EB026285 to M.B. and T32AI07328 to B.M.B.). PK is the beneficiary of a L’Oreal Poland and the Polish Ministry of Science and Higher Education and START Foundation for Polish Science scholarships. The Drag laboratory is supported by the “TEAM/2017-4/32” project, which is carried out within the TEAM program of the Foundation for Polish Science co-financed by the European Union under the European Regional Development Fund. The content is solely the responsibility of the authors and does not necessarily represent the official views of the National Institutes of Health.

