## Supplemental Methods, Figures and Tables for "Leveraging peptide substrate libraries to design inhibitors of bacterial Lon protease"

**Table of Contents:**

Supplemental Methods

Supplemental Synthesis Methods

Compound Characterization

Supplemental References

Supplemental Figures

Supplemental Tables

### Supplemental Methods

#### Lon purification

All purification steps following lysis were performed on ice or at 4 °C. Wild-type *Escherichia coli* Lon was purified as previously reported[1] with modifications. *E. coli* BL21 were transformed with pBAD33-lon[2], grown overnight in 50 ml 2xYT with 50 μg/ml chloramphenicol. The overnight culture was added to 4 l of 2xYT with 50 μg/ml chloramphenicol and grown to an $OD_{600}$ of approximately 0.8. Lon expression was induced by addition of L-arabinose (0.2%, w/v) and expression was allowed to proceed for 4 h. Cells were pelleted and frozen at -20 °C. The cell pellet was resuspended in 75 ml Lon lysis buffer (50 mM potassium phosphate pH 6.5, 2 mM DTT, and 10% glycerol) and lysed by incubation with 0.5 mg/ml lysozyme for 30 min on ice followed by probe sonication. The lysate was incubated with 0.1 mg/ml DNase I (Sigma-Aldrich) for 30 min on ice and then clarified by centrifugation. The lysate was further clarified by passage through a 0.2 μm vacuum filter before addition to the cellulose phosphate column.

Cellulose phosphate (Sigma-Aldrich) was prepared by washing 25 g with: 2 l water, 8 times; 500 ml 0.2 M NaOH, once; 2 l water, once; 500 ml 0.2 M HCl, once; and 2 l water. Washed cellulose phosphate was stored at 4°C as a 75% slurry in 100 mM potassium phosphate, pH 6.5 with 0.05% w/v of sodium azide. Cellulose phosphate was packed into a 50-ml column (GE Healthcare) for FPLC purification.

The lysate was applied to the cellulose phosphate column, washed with 100 ml of Lon lysis buffer, eluted with a 100 ml gradient from Lon lysis buffer to Lon elution buffer (400 mM potassium phosphate pH 6.5, 2 mM DTT, and 10% glycerol), and washed with 100 ml of Lon elution buffer. Elution fractions containing Lon were pooled and concentrated (Amicon Ultra 100KDa, EMD Millipore), loaded onto a Sephacryl S-300 column (GE Healthcare), and eluted in SEC buffer (25 mM Tris, pH 8, 100 mM KCl, 20% glycerol, 1 mM DTT). Elution fractions were tested for protease activity with FITC-casein (Sigma-Aldrich) and checked for purity by SDS-PAGE. Total protein (100 mg) was aliquoted and stored at -80°C.

**Inhibitor screens**

High throughput screens were run in black 384 well-plates. Each well contained 25 µl 2x Lon assay buffer (50 mM Tris, pH 8.0, 200 mM KCl, and 20 mM $MgCl_2$), 0.5 µl 4 µM Lon stock (40 nM hexamer final concentration), 0.5 µl 100 mM ATP (1 mM final concentration), 13.5 µl water, and 0.5 µl test compound in DMSO (1% final DMSO concentration). Compounds were incubated with Lon for 15 min at 37 °C before addition of 10 µl of 250 µM substrate **5** (50 µM final concentration) for a total assay volume of 50 µl. Fluorescence (ex. 360 nm, em. 460 nm) was measured every minute for 1 h at 37 °C in a microplate reader (BioTek Cytation 3). Each plate contained two columns of DMSO controls and two columns of 10 µM **BZ** for a total of 32 negative and positive control wells. Z' values for each plate ranged from 0.46 to 0.61. Initial cleavage rates were obtained by taking the slope of the initial timepoints of each curve and were normalized to DMSO-treated controls to yield percent activity.

**Fitting for substrate parameters**

Fluorescence measurements were converted to product concentration using a standard curve for 7-amino-4-carbamoylmethylcoumarin. Initial cleavage rates (v) were obtained by taking the slope of the linear portion of each curve and kinetic parameters were determined by performing nonlinear fits to the Michaelis-Menten equation (Equation S1).

$$v = \frac{k_{cat} C_E C_S}{K_M + C_S}$$

**Equation S1.** Michaelis-Menten equation for substrate parameters with v, initial cleavage rate; $k_{cat}$, catalytic rate; $C_E$, enzyme concentration; $C_S$, substrate concentration; and $K_M$, Michaelis constant

#### Fitting for inhibitor parameters

Initial cleavage rates were obtained by taking the slope of the initial timepoints of each curve and were normalized to DMSO-treated controls to yield percent activity. Percent activities were fit to inhibitor concentrations using a four-parameter logistic equation (Equation S2) to calculate $IC_{50}$ values.

$$A = A_{max} + \frac{A_{min} - A_{max}}{1 + \left(\frac{C_I}{IC_{50}}\right)^H}$$

**Equation S2.** Four parameter logistic regression model with A, percent activity; $C_I$, concentration of inhibitor; $A_{max}$, percent activity maximum dose; $A_{min}$, percent activity with no compound; $IC_{50}$; and H, Hill's slope.

Inactivation was assumed to proceed via a two-step inactivation mechanism[3]. Inactivation curves were fit to equation S3 to determine the observed inactivation rate constant ($k_{obs}$), which was then fit to Equation S4 to determine inactivation rates ($k_{inact}$) and inhibitor affinities ($K_I$).

$$C_P = \frac{v_i}{k_{obs}}[1 - exp(-k_{obs}t)$$

**Equation S3.** Nonlinear inactivation curve with $C_P$, concentration of product; $v_i$, initial rate; $k_{obs}$, observed inactivation rate; and t, time.

$$k_{obs} = \frac{k_{inact}C_I}{K_I + C_I}$$

**Equation S4.** Time-dependent inactivation curve with $k_{obs}$, observed inactivation rate; $k_{inact}$, inactivation rate; $C_I$, concentration of inhibitor; and $K_I$, inhibitor affinity.

#### Bacterial strains and plasmids

Antibiotics were used at the following concentrations: 50 µg/ml kanamycin and 50 µg/ml spectinomycin. Complete list of strains and plasmids is found in Table S2. pBAD33.lon was a gift from Robert Sauer (Addgene plasmid 22145). pCas and pTargetF were gifts from Sheng Yang (Addgene plasmids 62225 and 62226). Wild-type (MG1655) and the *sulA* knockout (JW0941-1; *sulA773(del)::kan*) *E. coli* were obtained from The Coli Genetic Stock Center (Yale University).

**Generation of *lon* deletion strain**

All enzymes were purchased from New England Biolabs. All PCR steps used Phusion polymerase. The deletion mutant was generated following procedures described by Jiang, et al[4]. The targeting plasmid, containing a sgRNA complementary to a region within the *lon* gene, was generated by amplifying pTargetF with primers pTargetF.rev/pTarget_lon.for, treating the PCR product with polynucleotide kinase and ligating the blunt ends to generate pTarget_lon. Homologous fragments flanking the *lon* gene (~500 bp fragments A and B in Figure S5a) were amplified from the chromosome using lon_us.for/lon_us.rev and lon_ds.for/lon_ds.rev. PCR products were annealed via 15 rounds of overlap extension and then the entire 1 kb sequence was amplified for 15 rounds using lon_us.for/lon_ds.rev. MG1655 harboring pCas was grown in LB with kanamycin at 30 °C. Cas9 expression was induced by incubation with 0.2% arabinose for 3 h and cells were made electrocompetent by washing and concentration in glycerol. Cells were electroporated with 100 ng pTarget_lon and 400 ng of the linear homologous fragments. Transformants were selected on LB plates with kanamycin and spectinomycin at 30 °C. Colonies were checked for *lon* deletion by PCR with lon_us_seq.for/lon_ds_seq.rev (Figure S5b). Cells were cured of pTarget_lon by culturing in LB with kanamycin and 0.5 mM IPTG overnight at 30 °C and then plating onto LB plates with kanamycin. Cells were cured of pCas by growing in LB at 37 °C.

### Supplemental Synthesis Methods

#### Solid-phase synthesis of peptide 7-amino-carbamoylmethylcoumarin (ACC) substrates 1-5

Fmoc-7-amino-4-carbamoylmethylcoumarin was synthesized and loaded onto Rink amide resin as previously described[5] to generate ACC-resin. ACC-resin (200 mg, 0.09 mmol ACC) was loaded into a synthesis cartridge and deprotected with 20% (v/v) piperidine in DMF for 30 min at room temperature. The resin was washed three times with DCM and three times with DMF. Fmoc-L-Phe-OH (174 mg, 0.45 mmol) was activated with HATU (171 mg, 0.45 mmol) and collidine (54.5 mg, 0.45 mmol) in DMF for 5 min and then added to the resin in DMF and allowed to react 16 h at room temperature with rotation. The resin was washed five times with DCM and five times with DMF. Any remaining free amines were acetylated by incubation of the resin with 20% (v/v) acetic anhydride in pyridine for 2 h at room temperature. The resin was washed five times with DCM, five times with DMF, twice with MeOH, and dried under vacuum.

Peptide substrates were synthesized using an automated peptide synthesizer (Biotage). The resin was washed five times with DMF after each deprotection and coupling step. For each substrate, 10 mg of Fmoc-Phe-ACC resin was loaded into a reactor. Resin was deprotected with 20% (v/v) piperidine in DMF for 30 min at room temperature. The amino acid to be coupled (5 eq., 0.025 mmol) was activated with HATU (9.5 mg, 0.025 mmol) and collidine (3 mg, 0.025 mmol) for five minutes, added to the resin, and allowed to react for 3 h. Three cycles of deprotection and coupling were used to add P2 (Fmoc-Leu-OH or Fmoc-hArg(Pbf)-OH) and P3 amino acids (Fmoc-Leu-OH, Fmoc-Lys(TFA)-OH, Fmoc-His(3-BOM)-OH, Fmoc-dhTrp(Boc)-OH, or Fmoc-nptGly-OH) and a 3-morpholine carboxylic acid cap to the resin. Substrates were cleaved from the resin and Boc- and Pbf-deprotected by addition of 95.0:2.5:2.5 TFA: $H_2O$:TIPS (twice for 1 h each). Substrates were precipitated by addition of diethyl ether (-20°C) to the cleavage solution, washed once with diethyl ether (-20°C), and dried under vacuum. The following products were obtained as powders.

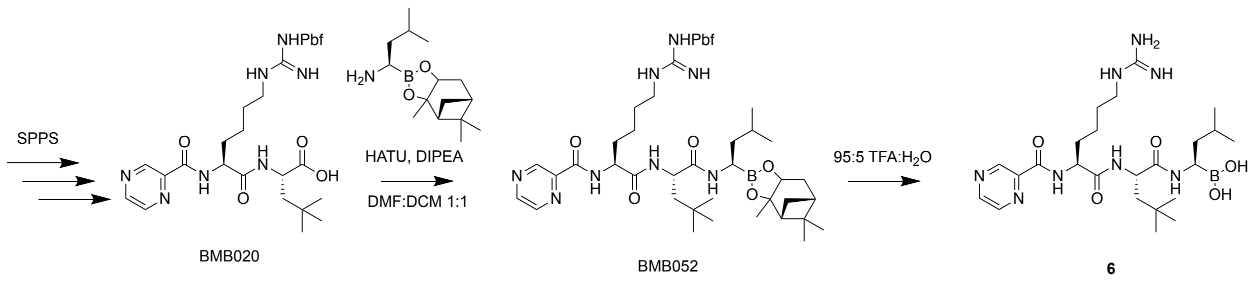

**Scheme S1.** Synthesis of **6**.

**H-hArg(Pbf)-nptGly-chlorotrityl resin (BMB019)**

Chlorotrityl resin (385 mg, 0.5 mmol) was washed twice with DCM and twice with DMF. After each coupling and deprotection, the resin was washed three time with DCM and three times with DMF. Fmoc-nptGly-OH (550 mg, 1.5 mmol) was activated with HATU (570 mg, 1.5 mmol) and DIEA (349 μl, 2.0 mmol), added to the resin, and allowed to react for 3 h. The resin was deprotected with 20% (v/v) piperidine in DMF for 30 min. Fmoc-hArg(Pbf)-OH (994 mg, 1.5 mmol) was activated with HATU (570 mg, 1.5 mmol) and DIEA (349 μl, 2.0 mmol), added to the resin, and allowed to react for 3 h. The resin was deprotected with 20% (v/v) piperidine/DMF for 30 min.

**Pyz-hArg(Pbf)-nptGly-OH (BMB020)**

Pyrazine carboxylic acid (46.5 mg, 0.375 mmol) was activated with HATU (143 mg, 0.375 mmol) and DIEA (79 μl, 0.45 mmol), added to BMB019 (200 mg, 0.15 mmol), and allowed to react for 3 h. The resin was washed five times with DCM and five times with DMF. The peptide was cleaved by treating the resin with 4:1 HFIP:DCM twice for 30 min each and then the resin was washed with DCM. The cleavage solution was dried under vacuum. The oil was dissolved in 50% (v/v) ACN/$H_2O$ and the product was purified by reverse-phase HPLC (50-95% ACN/$H_2O$ gradient on a $C_{18}$ column) and lyophilized to yield powder (79.7 mg, 0.118 mmol, 79% yield). LC-MS analysis found $[M+H]^+$ = 674.3; calculated 674.33. $^1H$ NMR (400 MHz, Chloroform-d) δ 9.34 (s, 1H), 8.76 (s, 1H), 8.56 (s, 1H), 4.72 (s, 1H), 4.59 (s, 1H), 3.24 (s, 2H), 2.97 (s, 2H), 2.90 (s, 1H), 2.52 (s,

3H), 2.47 (s, 3H), 2.10 (s, 3H), 2.07 – 1.51 (m, 6H), 1.47 (s, 6H), 1.33 – 1.18 (m, 1H), 0.92 (s, 9H).

**Pyz-hArg(Pbf)-nptGly-Leu-pinanediol boronate ester (BMB052)**

BMB020 (20.3 mg, 0.03 mmol) was dissolved in 1:1 DCM:DMF, activated with HATU (12.5 mg, 0.033 mmol) and DIEA (11.6 mg, 0.09 mmol) and then added to (R)-BoroLeu-(+)-Pinanediol trifluoroacetate (12.5 mg, 0.033 mmol) and allowed to react for 24 h. The reaction was dried under vacuum. The oil was resuspended in 60% (v/v) ACN/$H_2O$ and the product was purified by reverse-phase HPLC (40-95% ACN/$H_2O$ gradient on a $C_{18}$ column) and lyophilized to yield powder (5.9 mg, 0.0064 mmol, 21% yield). LC-MS analysis found $[M+H]^+$ = 921.6; calculated 921.54. $^1H$ NMR (400 MHz, Chloroform-d) δ 9.33 (s, 1H), 8.76 (s, 1H), 8.56 (s, 1H), 4.34 – 4.23 (m, 1H), 3.77 – 3.66 (m, 2H), 3.14 (dd, J = 7.5, 4.2 Hz, 3H), 2.98 (s, 2H), 2.55 (s, 3H), 2.50 (s, 3H), 2.11 (s, 3H), 1.50 – 1.18 (m, 29H), 0.97 – 0.85 (m, 15H), 0.84 – 0.81 (m, 3H).

**Pyz-hArg-nptGly-Leu-boronic acid (6)**

BMB052 (5.9 mg, 0.0064 mmol) was dissolved in 95:5 TFA:$H_2O$ and allowed to react for 2 h at room temperature. The reaction was dried under vacuum, resuspended in 30% (v/v) ACN/$H_2O$. The product was purified by reverse-phase HPLC (35-75% ACN/$H_2O$ gradient on a $C_{18}$ column) and lyophilized to yield powder (3.1 mg, 0.0058 mmol, 91% yield). LC-MS analysis found $[M+H]^+$ = 535.4; calculated 535.35. $^1H$ NMR (300 MHz, Methanol-d4) δ 9.21 (d, J = 1.5 Hz, 1H), 8.80 (d, J = 2.4 Hz, 1H), 8.69 (dd, J = 2.5, 1.5 Hz, 1H), 4.73 – 4.62 (m, 1H), 4.61 – 4.51 (m, 1H), 3.16 (t, J = 6.8 Hz, 2H), 2.72 (dd, J = 9.0, 6.2 Hz, 1H), 1.87 (tt, J = 14.0, 7.1 Hz, 2H), 1.77 – 1.56 (m, 6H), 1.55 – 1.40 (m, 2H), 1.37 – 1.22 (m, 1H), 0.95 (s, 9H), 0.93 (d, J = 2.4 Hz, 3H), 0.91 (d, J = 2.5 Hz, 3H).

### Compound Characterization

### Mo-Leu-Leu-Phe-7-amino-carbamoylmethylcoumarin (1)

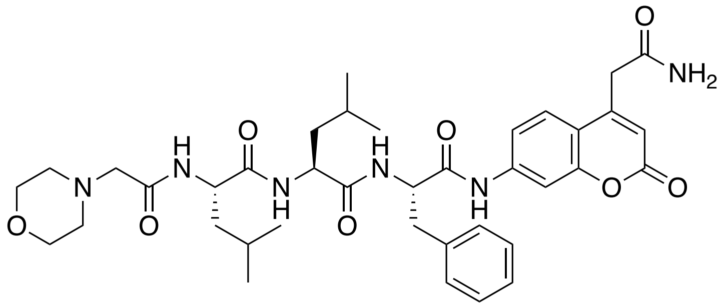

LC-MS analysis found $[M+H]^+$ = 719.4 ; calculated 719.37.

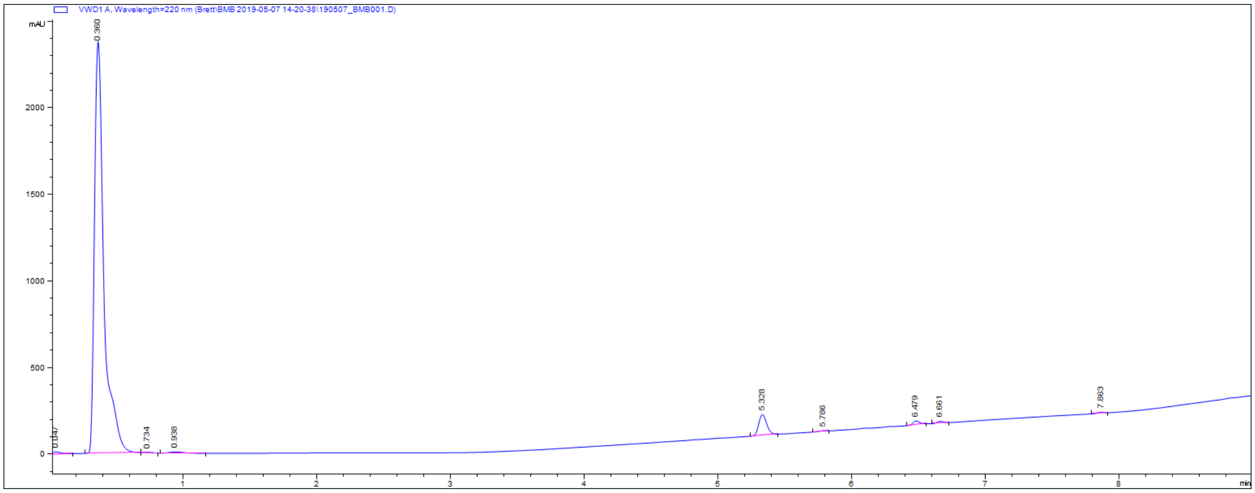


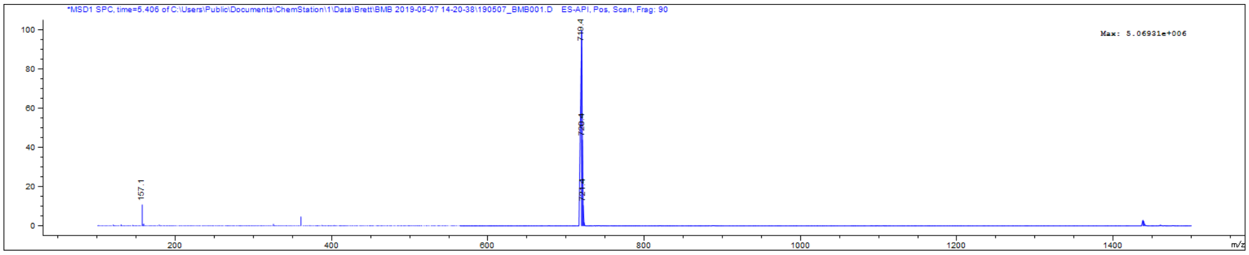

**Mo-hArg-Lys(TFA)-Phe-7-amino-carbamoylmethylcoumarin (2)**

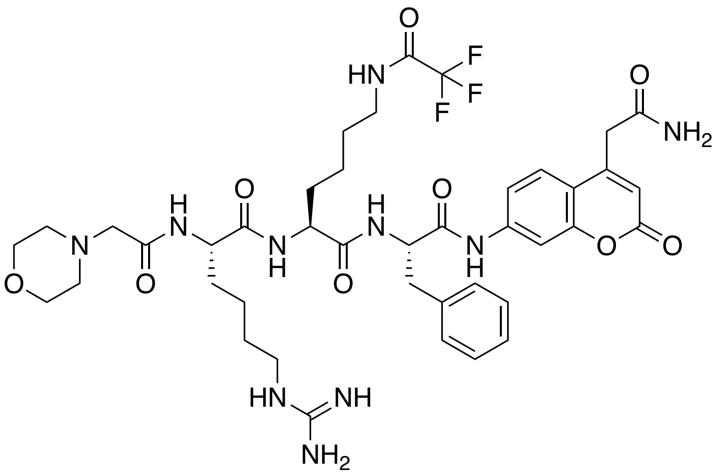

LC-MS analysis found $[M+H]^+$ = 887.5; calculated 887.39.

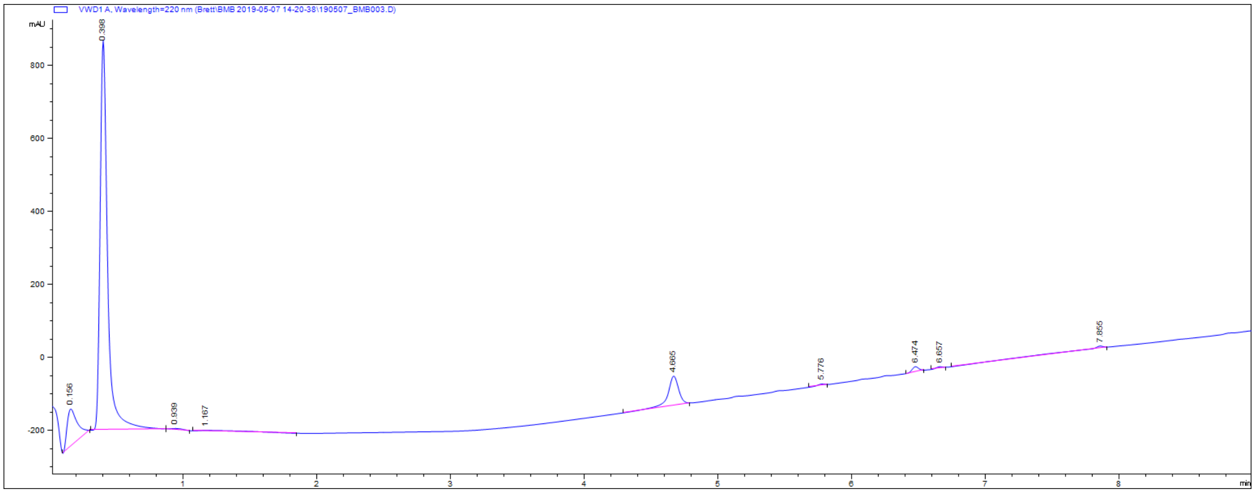


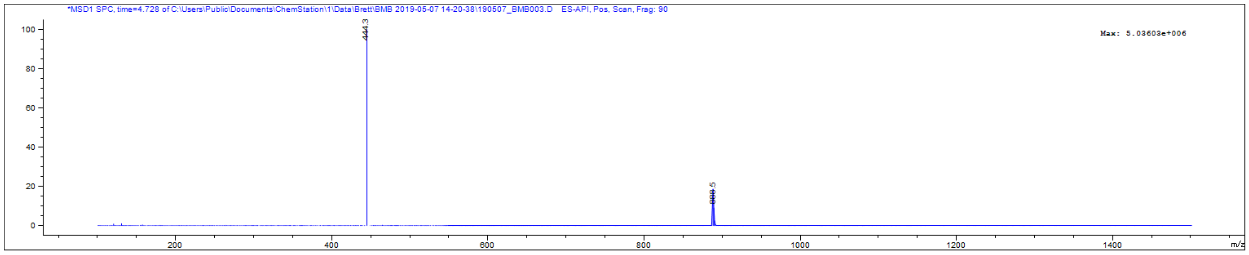

**Mo-hArg-His(3-BOM)-Phe-7-amino-carbamoylmethylcoumarin (3)**

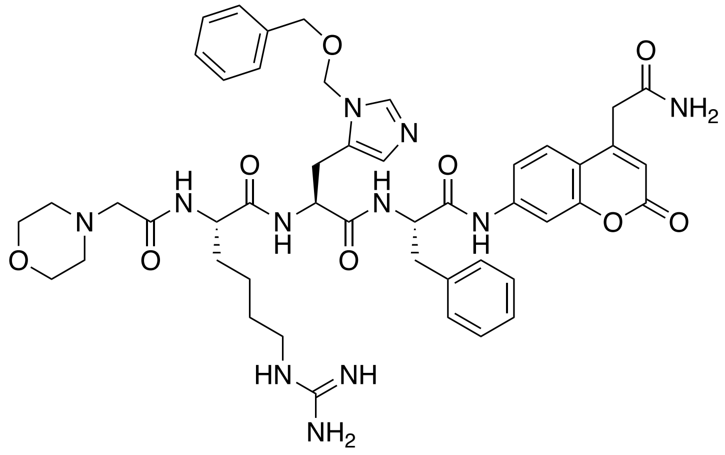


LC-MS analysis found $[M+H]^+$ = 920.5; calculated 920.43.

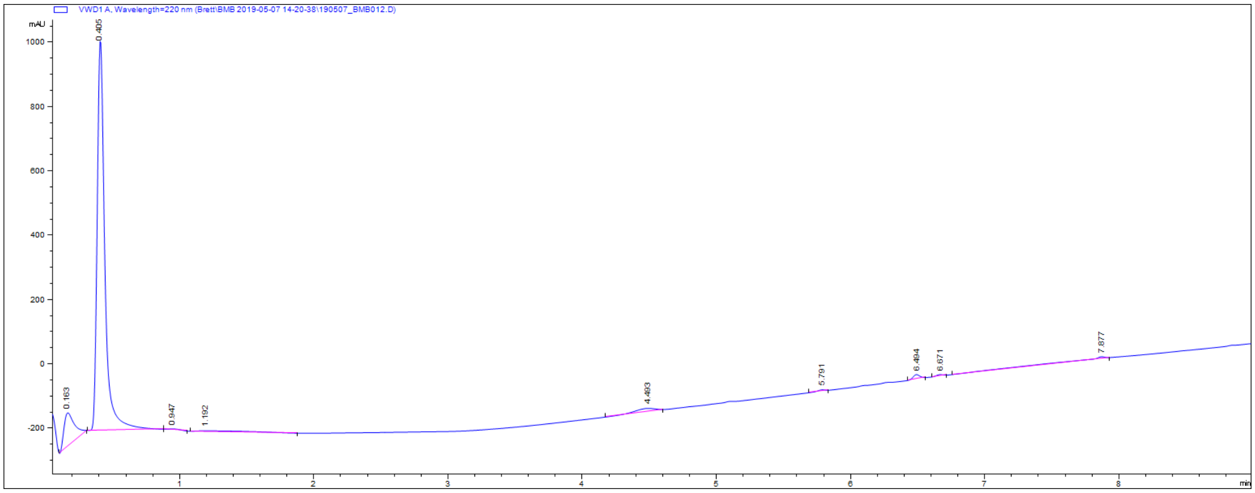


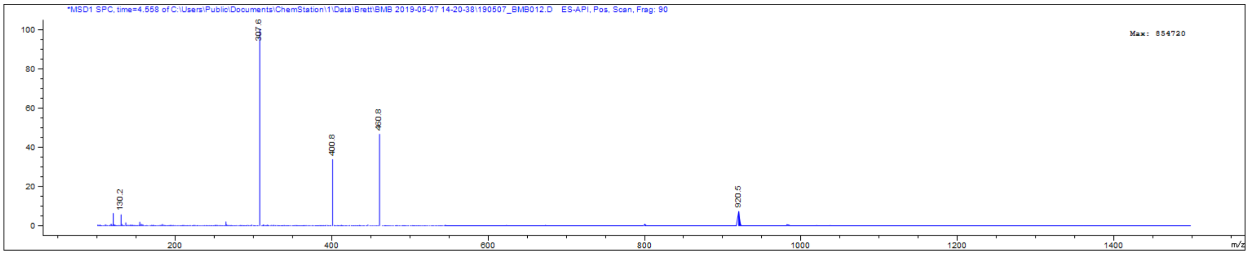

**Mo-hArg-dhTrp-Phe-7-amino-carbamoylmethylcoumarin (4)**

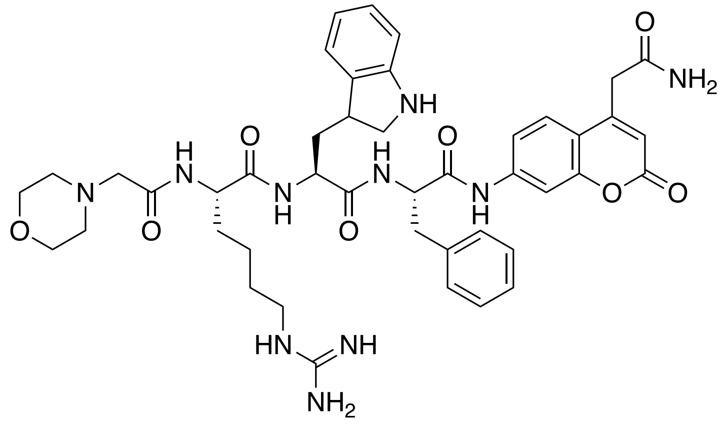


LC-MS analysis found $[M+H]^+$ = 851.5; calculated 851.41.

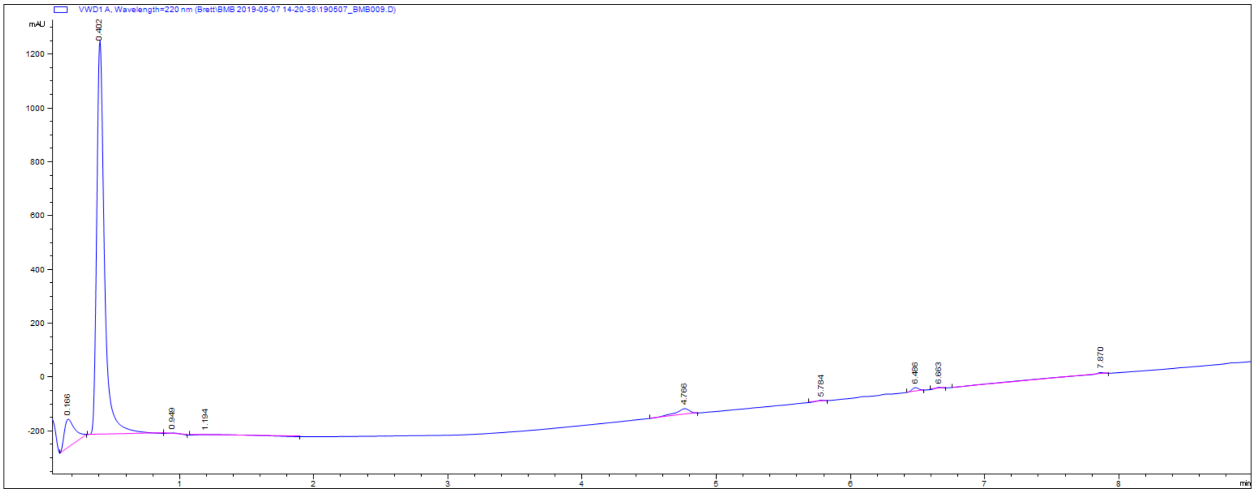


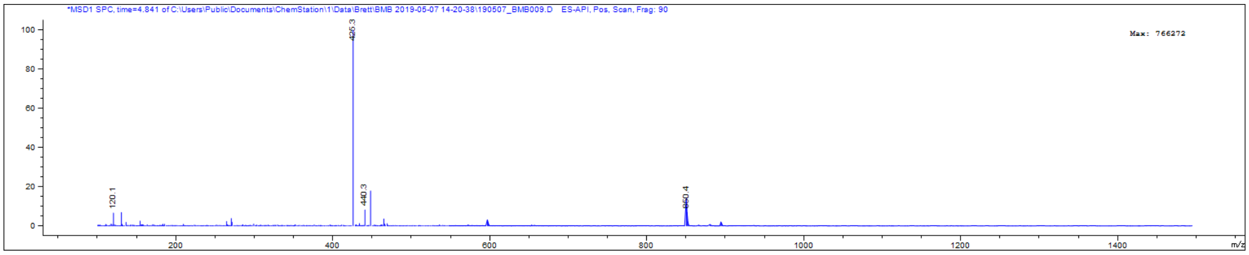

**Mo-hArg-nptGly-Phe-7-amino-carbamoylmethylcoumarin (5)**

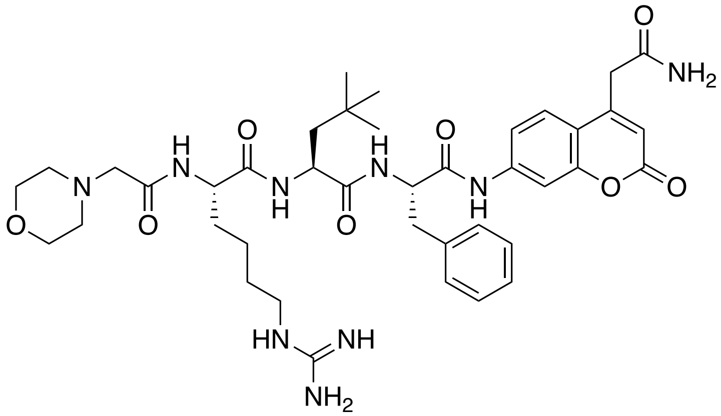


LC-MS analysis found $[M+H]^+$ = 790.5; calculated 790.42.

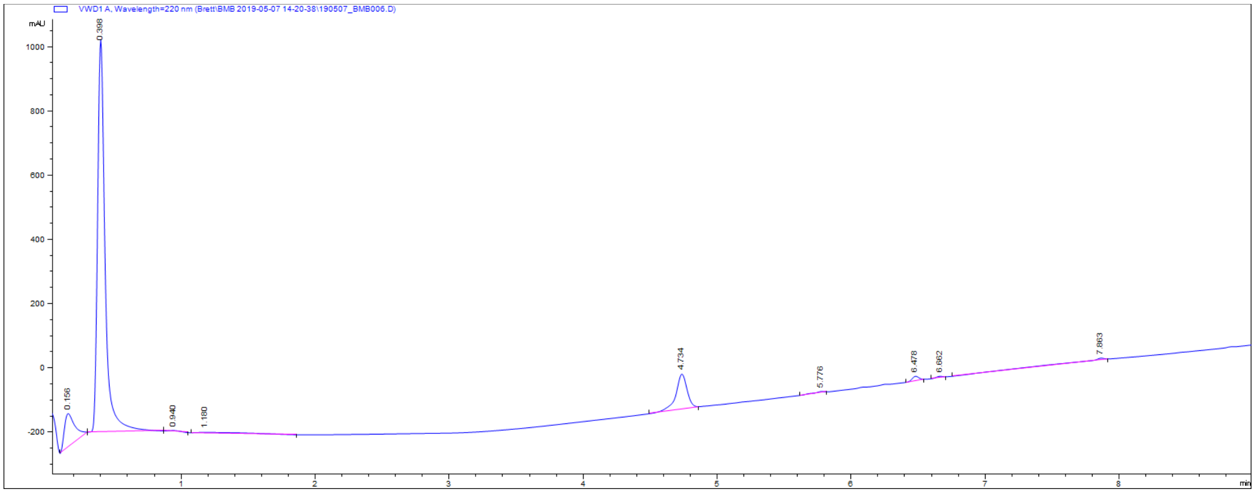


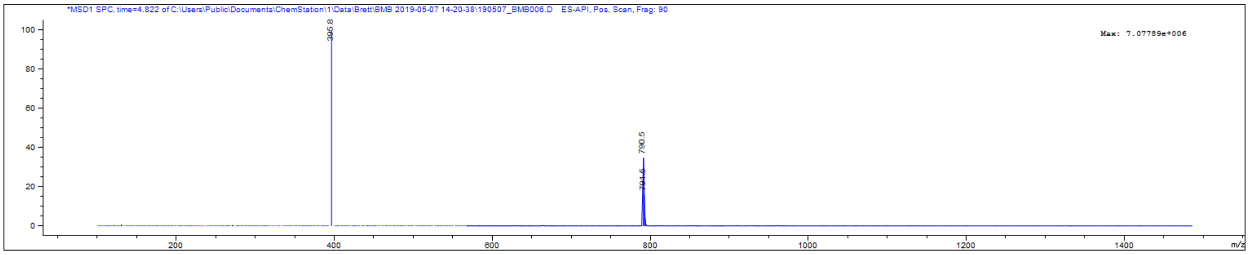

### Pyz-hArg(Pbf)-nptGly-OH (BMB020)

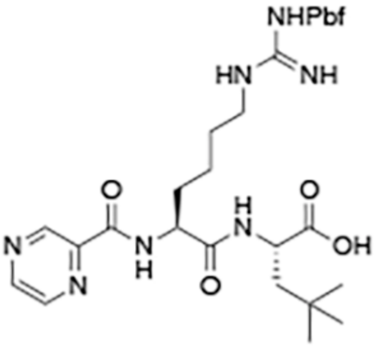


BMB020

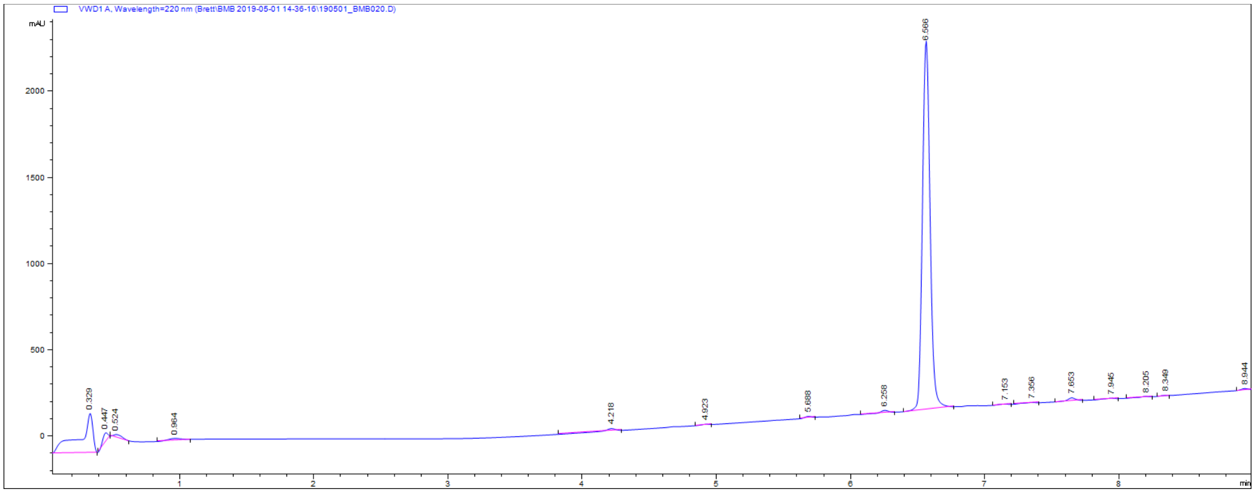


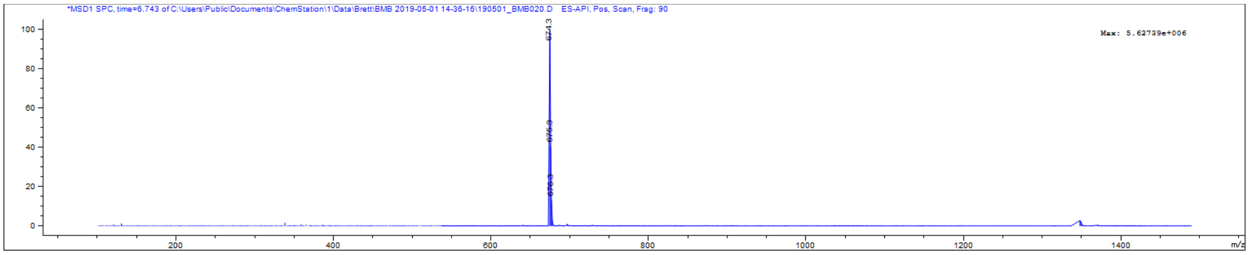

### Pyz-hArg(Pbf)-nptGly-Leu-pinanediol boronate ester (BMB052)

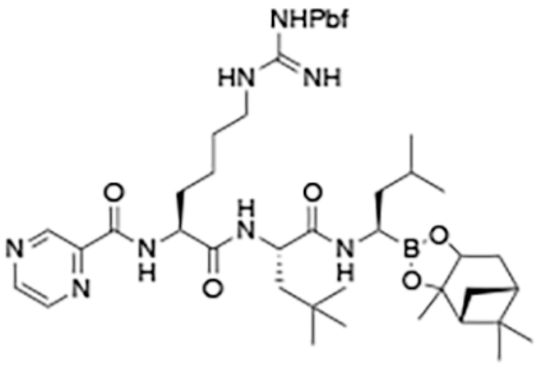


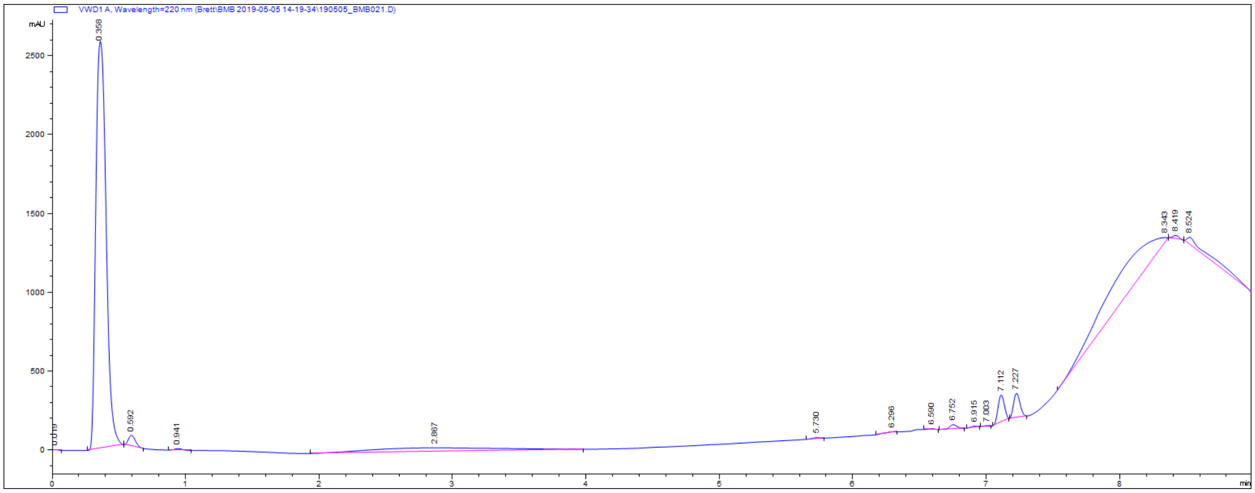


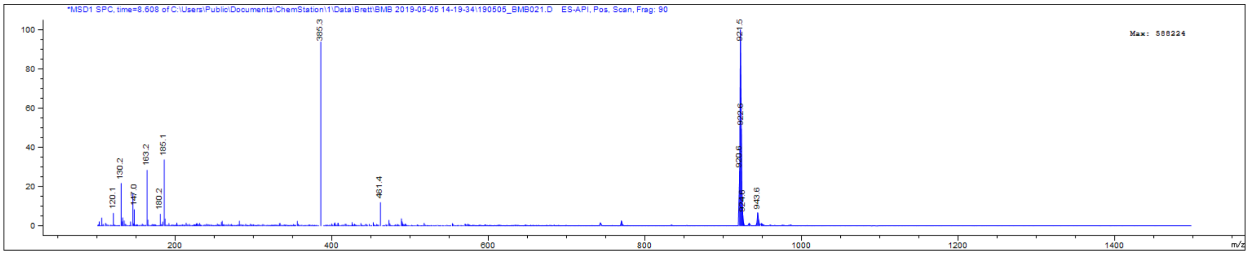

### Pyz-hArg-nptGly-Leu-boronic acid (6)

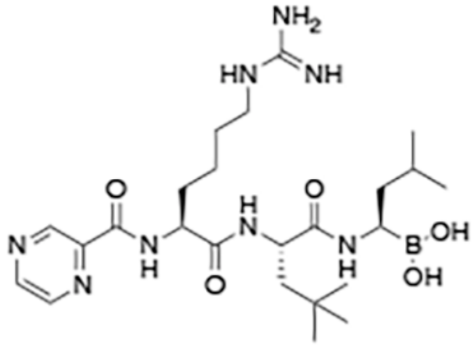


**6**

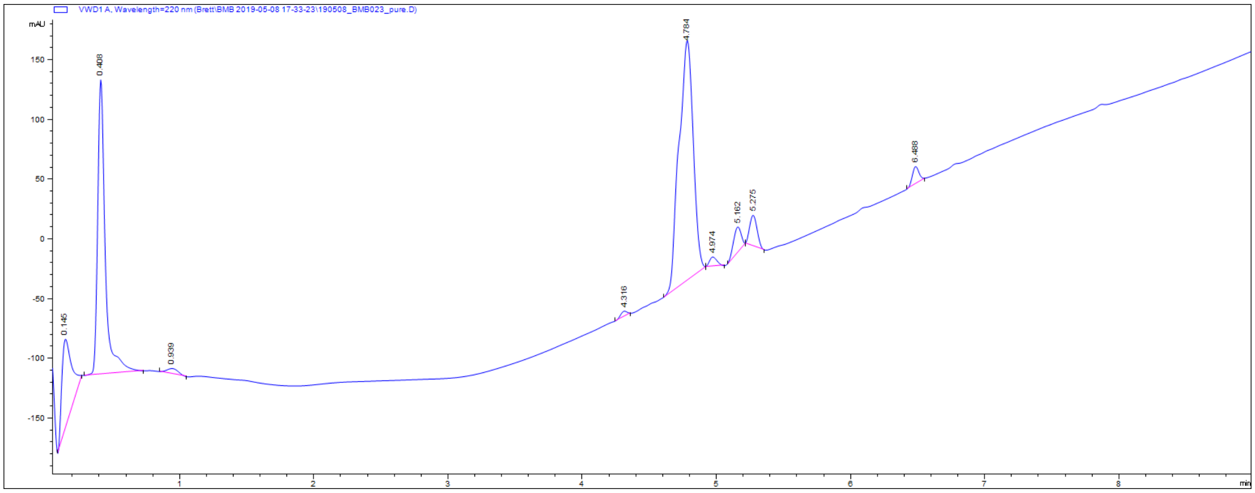


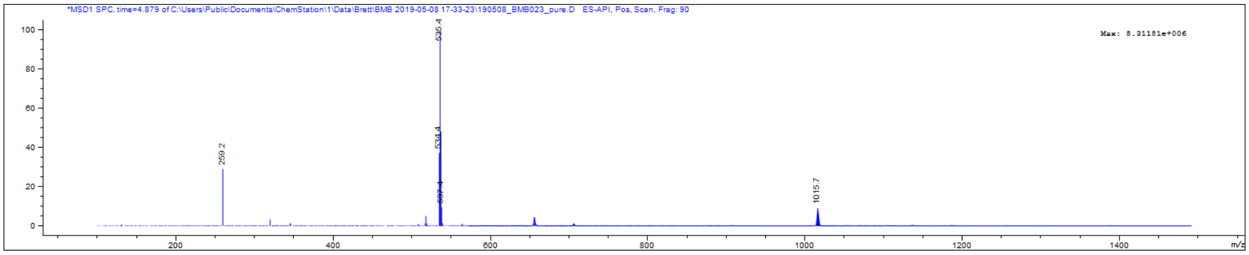

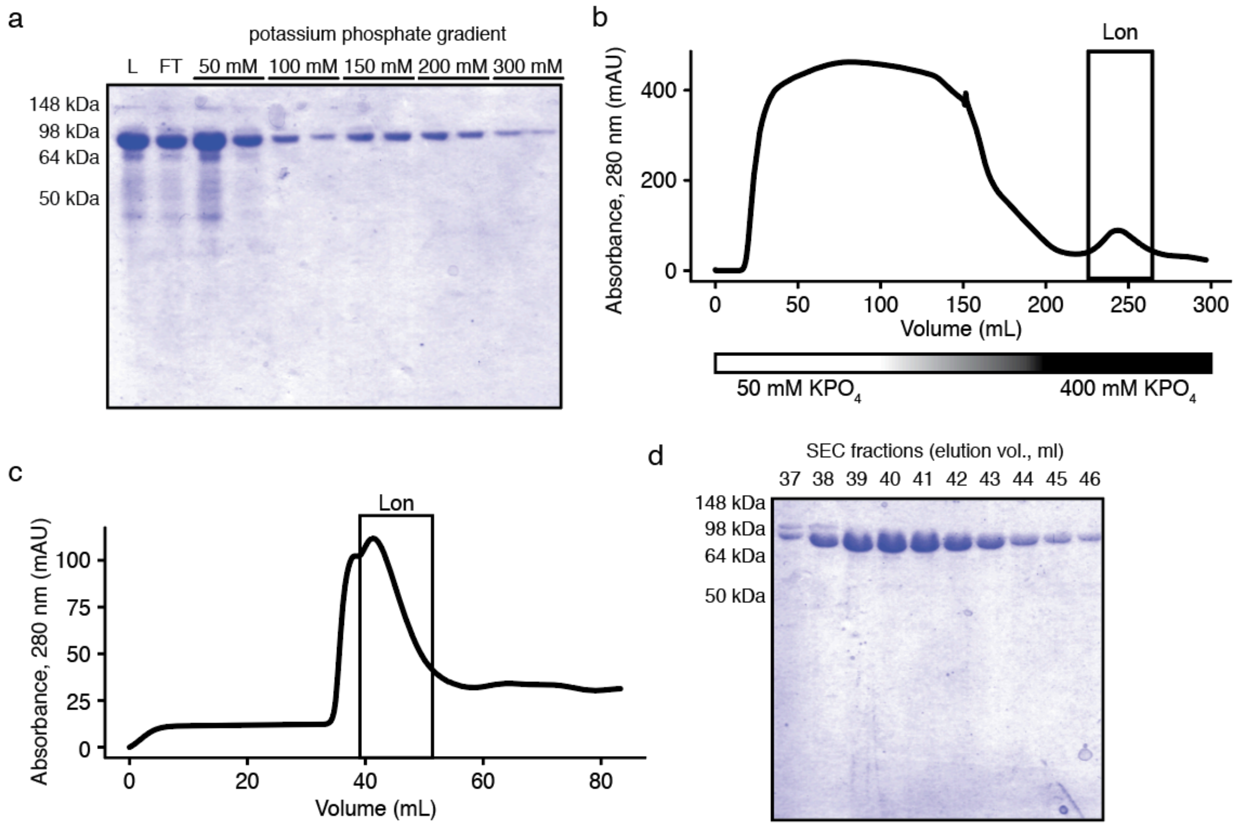


**Figure S1. Purification of Lon.** (a) SDS-PAGE analysis of cellulose phosphate purification. L, lysate; FT, flow-through; and numbers indicate fractions eluted with each potassium phosphate concentration. (b) UV absorbance trace of FPLC purification on the cellulose phosphate column. Potassium phosphate concentration is indicated below. The peak containing Lon is indicated. (c) UV absorbance trace of FPLC purification with Sephacryl S-300 column. The peak containing active Lon is indicated. (d) SDS-PAGE analysis of size exclusion fractions. Fractions 39-46 were pooled. Gels were stained with Coomassie.

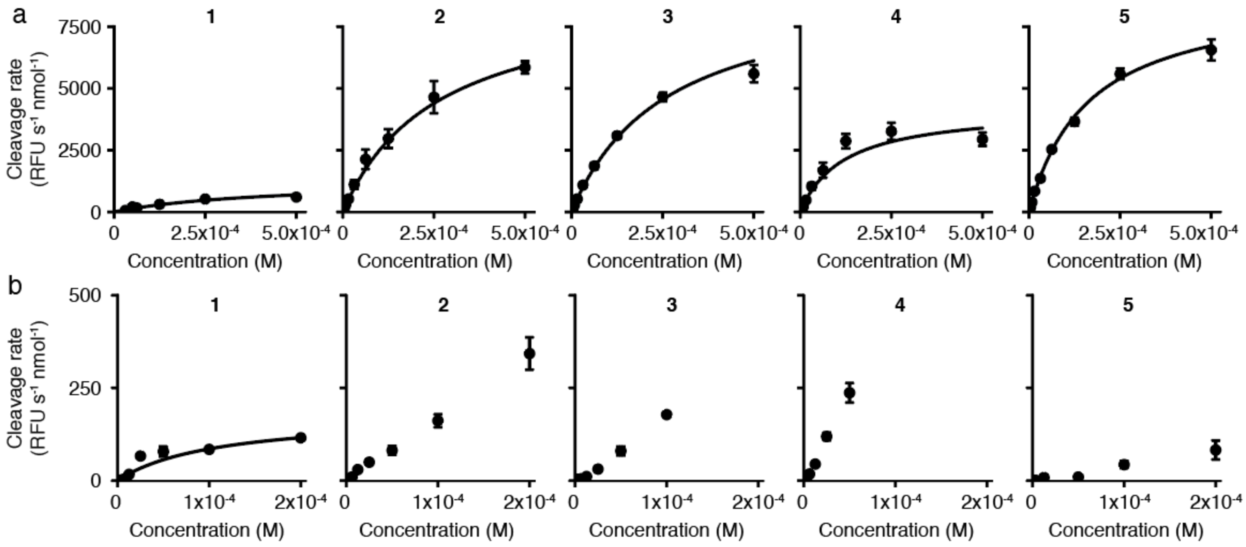


c

h20S kinetic parameters

| | P3 | P2 | $k_{cat}$ ($10^{-3}$ $s^{-1}$) | $K_M$ ($10^{-6}$ M) | $k_{cat}/K_M$ ($M^{-1}$ $s^{-1}$) |
|---|---|---|---|---|---|
| **1** | Leu | Leu | 5.7 ± 4.2 | 110 ± 30 | 52 ± 41 |
| **2** | hArg | Lys(TFA) | ND | ND | ND |
| **3** | hArg | His(3-BOM) | ND | ND | ND |
| **4** | hArg | dhTrp | ND | ND | ND |
| **5** | hArg | nptGly | ND | ND | ND |

**Figure S2. Kinetic analysis of fluorogenic substrates.** Initial cleavage rates vs. substrate concentration for (a) 40 nM Lon or (b) 2 nM h20S (mean ± standard deviation, n=3). Cleavage rates are normalized to the enzyme concentration. Fits to Equation S1 are indicated by solid black lines. (c) Kinetic parameters for cleavage by h20S (mean ± standard deviation, n=3). Cleavage of **2**–**5** by h20S did not saturate at the concentrations tested and parameters were not calculated.

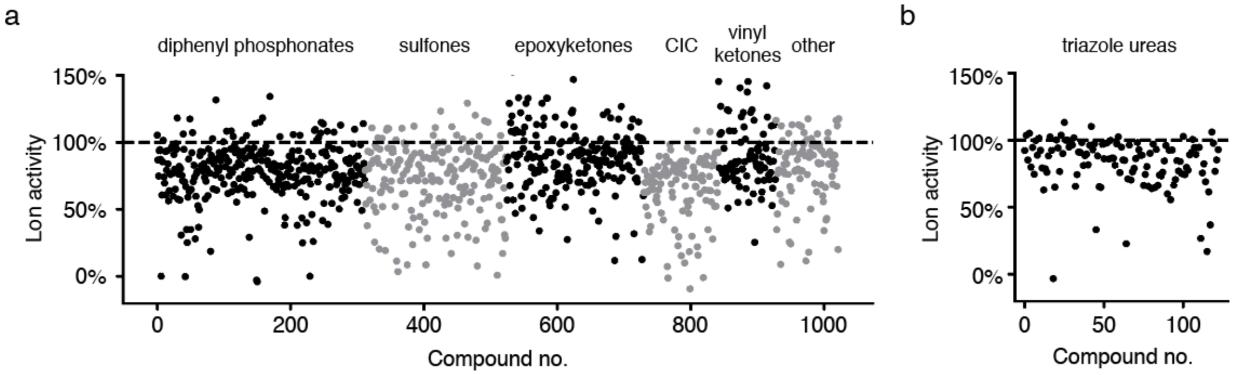


c

| Compound | Class | $IC_{50}$ ($10^{-6}$ M) |
|---|---|---|
| **BZ** | boronic acid | 0.50 |
| **WRR-000139** | vinyl sulfone | 17 |
| **JCP-000656** | epoxyketone | 22 |
| **JCP-000453** | vinyl sulfone | 28 |
| **JCP-000342** | chloroisocoumarin | 29 |
| **JCP-000469** | fluoromethyl ketone | 30 |
| **JCP-000247** | chloroisocoumarin | 39 |
| **WRR-000230** | vinyl sulfone | 42 |
| **WRR-000442** | vinyl sulfone | 46 |
| **JCP-000180** | chloroisocoumarin | 95 |
| **JCP-000126** | diphenyl phosphonate | 96 |

**Figure S3. Screens for Lon inhibitors.** Screening results for (a) 10 µM or (b) 1 µM of the indicated compound classes. Lon (40 nM) was incubated with each inhibitor for 15 min at 37 °C and then incubated with 50 µM **5** to measure activity. Results are normalized to DMSO controls. (c) $IC_{50}$ values for the best inhibitors from the screens and for **BZ**.

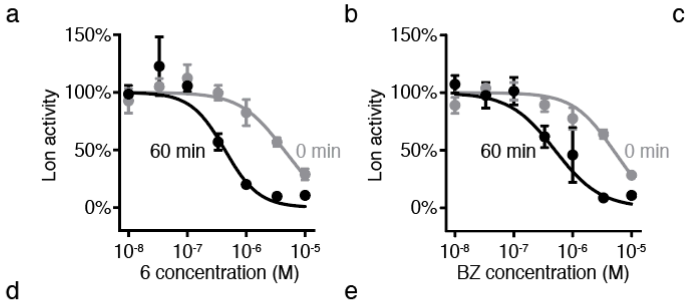


c

| | Pre-incubation (min) | $IC_{50}$ ($10^{-9}$ M) |
|---|---|---|
| 6 | 0 | 4400±380 |
| | 60 | 430±140 |
| BZ | 0 | 4900±600 |
| | 60 | 500±230 |

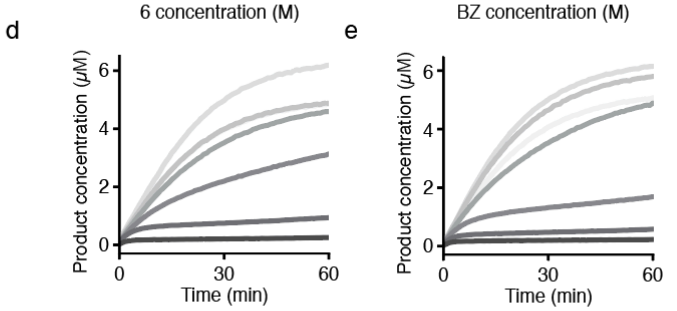


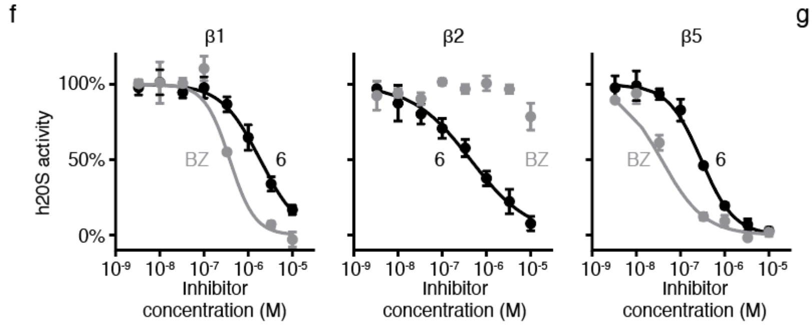


g

| Subunit | Substrate | Inhibitor | $IC_{50}$ ($10^{-9}$ M) |
|---|---|---|---|
| β1 | Z-LLE-AMC | 6 | 1900±140 |
| | | BZ | 380±20 |
| β2 | Boc-LRR-AMC | 6 | 430±40 |
| | | BZ | ND |
| β5 | Suc-LLVY-AMC | 6 | 290±6.4 |
| | | BZ | 37±7.1 |

**Figure S4. Kinetic analysis of inhibitors.** Inhibition of Lon as determined by initial cleavage rates by (a) **6** or (b) **BZ** following 60 min (black circles) or 0 min (gray circles) enzyme-inhibitor pre-incubation (mean ± standard deviation, n=3). Reactions contained 40 nM Lon (240 nM active sites) and 50 μM **5**. Fits to Equation S2 are indicated by solid lines. (c) $IC_{50}$ values for **6** and **BZ** against Lon (mean ± standard deviation, n=3). Reaction progress curves for inhibition of Lon by (d) **6** or (e) **BZ** with no pre-incubation. Lines represent the mean of three measurements and shading indicates inhibitor concentrations, ranging from 10 nM (light gray) to 10 μM (black) in half-log increments. (f) Inhibition of h20S subunits as determined by initial cleavage rates by **6** (black circles) or **BZ** (gray circles) (mean ± standard deviation, n=3). Reactions contained 2 nM h20S, 24 nM PA28, and 50 μM substrate. Subunit activities were assayed separately using subunit-specific substrates: Z-LLE-AMC, Boc-LRR-AMC, and Suc-LLVY-AMC specific for the β1, β2, and β5 subunits, respectively. (g) $IC_{50}$ values for **6** and **BZ** against h20S subunits (mean ± standard deviation, n=3).

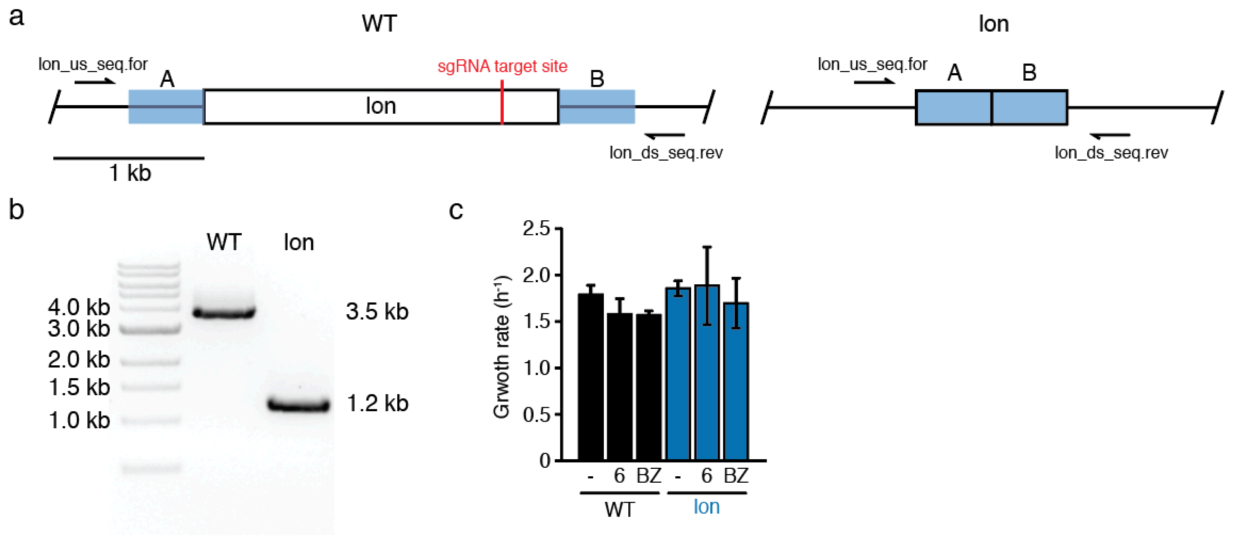


**Figure S5. Generation of the *lon* mutant.** (a) *E. coli* chromosome for wild-type (left) and *lon* mutant (right). The targeted site for CRISPR counterselection is indicated by a red line. Homologous flanking regions are indicated in blue. Primers used to check deletion are shown. (b) Colony PCR products for wild-type and *lon* deletion strains (lon_us_seq.for/lon_ds_seq.rev). (c) Exponential growth rates for each strain treated with DMSO (-) or the indicated compounds at 100 µM in LB at 37 °C (mean ± standard deviation, n=4; unpaired t-test comparing each condition to wild-type cells treated with DMSO, p>0.05 for all conditions).

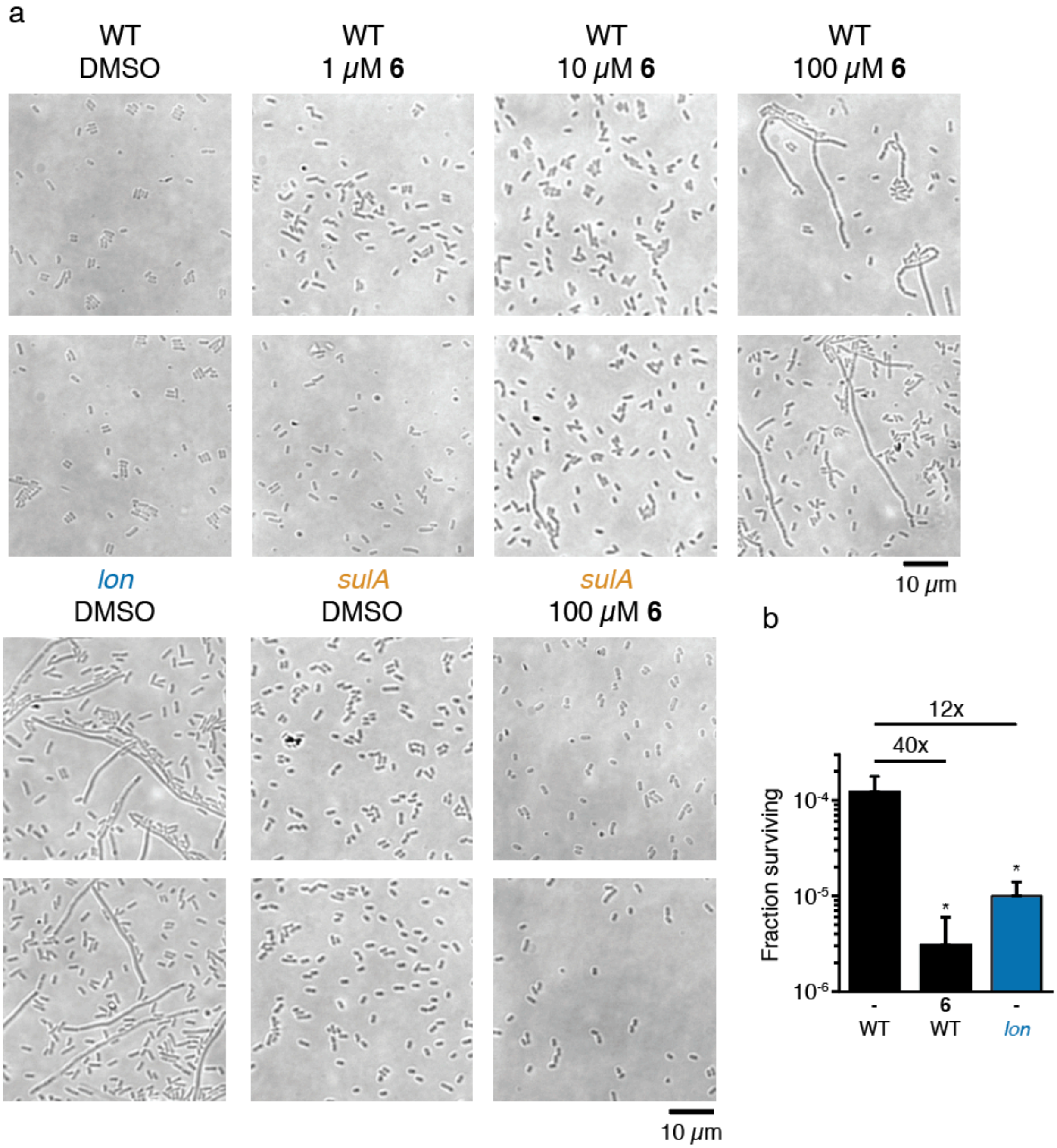


**Figure S6. Lon phenotypes.** (a) Additional microscopy images of conditions presented in Figure 4. (b) Fraction of *E. coli* cells surviving treatment with 10 μg/mL ciprofloxacin in M9. Cells were co-treated with ciprofloxacin and either DMSO (-) or 100 μM **6** for 4 h (mean ± standard deviation, n=3; unpaired t-tests comparing each sample to wild-type cells treated with DMSO: *, p<0.05). Fold reduction compared to wild-type cells treated with DMSO is indicated above.

**Table S1. HyCoSuL percent maximum cleavage rates for P2 and P3 positions.**

| Amino Acid Number. | Amino Acid | P2 Average Percent Activity | P3 Average Percent Activity | Amino Acid Number | Amino Acid | P2 Average Percent Activity | P3 Average Percent Activity |
|---|---|---|---|---|---|---|---|
| 1 | Ala | 0% | 15% | 62 | Abu | 15% | 0% |
| 2 | Arg | 28% | 10% | 63 | nptGly | 100% | 9% |
| 3 | Asn | 0% | 0% | 64 | Trp(Me) | 35% | 5% |
| 4 | Asp | 0% | 0% | 65 | Arg(Z)2 | 12% | 0% |
| 5 | Glu | 4% | 0% | 66 | 2-Aoc | 19% | 0% |
| 6 | Gln | 19% | 6% | 67 | Arg(NO2) | 51% | 0% |
| 7 | Gly | 0% | 0% | 68 | Glu(OMe) | 18% | 12% |
| 8 | His | 22% | 11% | 69 | Hnv | 24% | 21% |
| 9 | Ile | 0% | 0% | 70 | Phe(3F) | 5% | 0% |
| 10 | Leu | 42% | 13% | 71 | 4-Pal | 28% | 0% |
| 11 | Lys | 23% | 13% | 72 | Bip | 0% | 0% |
| 12 | Nle | 12% | 25% | 73 | Met(O) | 14% | 0% |
| 13 | Phe | 0% | 7% | 74 | His(Bzl) | 21% | 0% |
| 14 | Pro | 0% | 0% | 75 | Abu(Bht) | 15% | N.D. |
| 15 | Ser | 0% | 0% | 76 | Tyr(Bzl) | 0% | 0% |
| 16 | Thr | 0% | 0% | 77 | 2,3-dhAbu | 0% | 0% |
| 17 | Trp | 36% | 13% | 78 | hLeu | 26% | 19% |
| 18 | Tyr | 4% | 36% | 79 | Lys(Ac) | 40% | 28% |
| 19 | Val | 5% | 18% | 80 | Dap | 4% | 0% |
| 20 | D-Ala | 0% | 0% | 81 | Ala(Bth) | 5% | 0% |
| 21 | D-Arg | 0% | 0% | 82 | Hyp(Bzl) | 0% | 0% |
| 22 | D-Asn | 0% | 0% | 83 | dhTrp | 93% | 49% |
| 23 | D-Asp | 0% | 0% | 84 | Met(O)2 | 18% | 0% |
| 24 | D-Glu | 0% | 0% | 85 | hTyr | 56% | 28% |
| 25 | D-Gln | 0% | 0% | 86 | 1-Nal | 46% | 0% |
| 26 | D-His | 0% | 0% | 87 | Thz(Bzl) | 0% | 0% |
| 27 | D-Leu | 0% | N.D. | 88 | Phe(3,4-Cl2) | 0% | 0% |
| 28 | D-Lys | 0% | 0% | 89 | hArg | 64% | 100% |
| 29 | D-Phg | 0% | 0% | 90 | Lys(TFA) | 67% | N.D. |
| 30 | D-Phe | 0% | 0% | 91 | Thr(Bzl) | 0% | N.D. |
| 31 | D-Pro | 0% | 0% | 92 | Dab(Z) | 0% | N.D. |
| 32 | D-Ser | 0% | 0% | 93 | Chg | 6% | 0% |
| 33 | D-Thr | 0% | 0% | 94 | Asp(Bzl) | 3% | 0% |
| 34 | D-Trp | 0% | 0% | 95 | Nva | 15% | 20% |
| 35 | D-Tyr | 0% | 0% | 96 | Hyp | 0% | 0% |
| 36 | D-Val | 0% | N.D. | 97 | hCit | 41% | 14% |
| 37 | Tyr(Me) | 0% | 0% | 98 | Phe(4-Cl) | 0% | 0% |
| 38 | Phe(F5) | 0% | 0% | 99 | Phe(2-F) | 10% | 0% |
| 39 | Idc | 0% | 0% | 100 | Asp(OMe) | 7% | 0% |
| 40 | hTyr(Me) | 15% | 0% | 101 | Oic | 0% | 0% |
| 41 | Bpa | 5% | 4% | 102 | Ser(Ac) | 0% | 0% |
| 42 | Phe(3-Cl) | 0% | 0% | 103 | Glu(Bzl) | 19% | 0% |
| 43 | Phe(4-F) | 4% | 0% | 104 | Pip | 0% | 0% |
| 44 | Phe(2-Cl) | 16% | 0% | 105 | Phe(3,4-F2) | 8% | 0% |
| 45 | Phe(4-Me) | 0% | 9% | 106 | His(3-Bom) | 80% | 4% |
| 46 | 3-Pal | 28% | 0% | 107 | Agb | 22% | 0% |
| 47 | 2-Nal | 0% | 8% | 108 | Ser(Bzl) | 16% | 0% |
| 48 | Cha | 56% | 0% | 109 | Phe(4-I) | 0% | 8% |
| 49 | Tyr(2,6-Cl2-B | 0% | 10% | 110 | Orn | 16% | 6% |
| 50 | Ala(2-thienyl | 5% | 0% | 111 | Hse(Bzl) | 40% | 3% |
| 51 | hPhe | 10% | 0% | 112 | Agp | 8% | 0% |
| 52 | Asp(OChx) | 3% | 30% | 113 | 2-Igl | 16% | 0% |
| 53 | Tyr(2-Br-Z) | 16% | 0% | 114 | Aze | 0% | 0% |
| 54 | Phg | 8% | 0% | 115 | Bta | 9% | 10% |
| 55 | Tle | 4% | 0% | 116 | Hse | 13% | 0% |
| 56 | Lys(2Cl-Z) | 7% | 0% | 117 | dhLeu | 0% | 0% |
| 57 | Cys(4-MeBzl | 21% | 0% | 118 | Cys(Bzl) | 13% | N.D. |
| 58 | Phe(4-NH) | 9% | 0% | 119 | Dab | 3% | 0% |
| 59 | B-Ala | 0% | 0% | 120 | dhPro | 0% | 8% |
| 60 | Cit | 25% | 0% | 121 | Nle(O-Bzl) | 28% | 0% |
| 61 | Glu(OChx) | 44% | 0% | | | | |

**Table S2. List of bacterial strains and plasmids**

| Designation | Description | Source |
|---|---|---|
| **Strains** | | |
| WT | MG1655 | Yale Coli Genetic Stock Center |
| *lon* | MG1655 *Δlon* | This study |
| *sulA* | BW25113 *sulA773(del)::kan* | Yale Coli Genetic Stock Center; Baba et al.[6] |
| **Plasmids** | | |
| pBAD33.lon | Expression plasmid for Lon | Addgene; |
| pCas | For CRISPR-Cas mediated knockouts | Addgene; Jiang, et al. |
| pTargetF | For CRISPR-Cas mediated knockouts | Addgene; Jiang, et al. |
| pTargetF_lon | pTargetF with sgRNA for *lon* | This study |

**Table S3. List of primers**

| Designation | Sequence (5' to 3') |
|---|---|
| pTargetF.rev | ACTAGTATTATACCTAGGACTG |
| pTarget_lon.for | GTGCGCGTGCGGAAAAACTGGTTTTAGAGCTAGAAATAGC |
| lon_us.for | GCTCTGATTCAGATCCTC |
| lon_us.rev | GTGCATTTTGCGCTCTCTCTTAGTTTAATTTCCG |
| lon_ds.for | CTAAGAGAGAGCGCAAAATGCACTAATAAAAACAG |
| lon_ds.rev | CTTGAACTTCGTCACATCC |
| lon_us_seq.for | GAACCGGAAGATCTGATCAAG |
| lon_ds_seq.rev | CGCAGAAAGTCACCGAATAC |
